# The Layer 7 Cortical Interface: A Scalable and Minimally Invasive Brain–Computer Interface Platform

**DOI:** 10.1101/2022.01.02.474656

**Authors:** Mark Hettick, Elton Ho, Adam J. Poole, Manuel Monge, Demetrios Papageorgiou, Kazutaka Takahashi, Morgan LaMarca, Daniel Trietsch, Kyle Reed, Mark Murphy, Stephanie Rider, Kate R. Gelman, Yoon Woo Byun, Timothy Hanson, Vanessa Tolosa, Sang-Ho Lee, Sanjay Bhatia, Peter E. Konrad, Michael Mager, Craig H. Mermel, Benjamin I. Rapoport

## Abstract

Progress toward the development of brain–computer interfaces has signaled the potential to restore, replace, and augment lost or impaired neurological function in a variety of disease states. Existing approaches to developing high-bandwidth brain–computer interfaces rely on invasive surgical procedures or brain-penetrating electrodes, which limit addressable applications of the technology and the number of eligible patients. Here we describe a novel approach to constructing a neural interface, comprising conformable thin-film electrode arrays and a minimally invasive surgical delivery system that together facilitate bidirectional communication with large portions of the cortical surface (enabling both recording and stimulation). We demonstrate the feasibility and safety of delivering reversible implants containing over 2,000 microelectrodes to multiple functional regions in both hemispheres of the brain simultaneously, without requiring a craniotomy or damaging the cortical surface, at an effective insertion rate faster than 40 ms per channel. We further evaluate the performance of this system immediately following implantation for high-density neural recording and visualizing cortical surface activity at spatial and temporal resolutions and extents not previously possible in multiple preclinical large animal studies as well as in a five-patient pilot clinical study involving both anesthetized and awake neurosurgical patients. We characterize the spatial scales at which sensorimotor activity and speech are represented at the cortical surface, demonstrate accurate neural decoding of somatosensory, visual, and volitional walking activity, and achieve precise neuromodulation through cortical stimulation at sub-millimeter scales. The resulting system generates 90 Gb/h of electrophysiologic data, and demonstrates the highly scalable nature of micro-electrocorticography and its utility for next-generation brain-computer interfaces that may expand the patient population that could benefit from neural interface technology.

## 1 Introduction

Brain–computer interfaces have shown promise as systems for restoring, replacing, and augmenting lost or impaired neurological function in a variety of contexts, including paralysis from stroke and spinal cord injury, blindness, and some forms of cognitive impairment^1–21^. Multiple innovations over the past several decades have contributed to the potential of these neural interfaces, including advances in the areas of applied neuroscience and multichannel electrophysiology^22–31^, mathematical and computational approaches to neural decoding^27,32–38^, power-efficient custom electronics and the development of application-specific integrated circuits^39–51^, as well as materials science and device packaging^52–59^. Nevertheless, the practical impact of such systems remains limited, with only a small number of patients worldwide having received highly customized interfaces through clinical trials^60–62^.

In order to achieve meaningful clinical impact on the large populations of patients who stand to benefit from brain–computer interface technologies^63–68^, surgical procedures involved in implanting neural interfaces should be minimally invasive, reversible, and avoid damaging neural tissue. Advanced brain–computer interfaces require collection and processing of large amounts of neural data, potentially spanning multiple brain regions. As a result, high-density microelectrode arrays have been replacing more traditional macroelectrode arrays, offering smaller features and improved spatial resolution^69–74^. Systems for clinical use should also demonstrate a high degree of scalability in terms of channel count and speed of implantation. Tissue damage and total procedural time ideally should not increase in proportion to channel number, as contemporary channel counts, already reaching the many thousands, will likely increase by orders of magnitude with further progress in this field.

Microelectrode arrays that penetrate the brain have facilitated high-spatial-resolution recordings for brain–computer interfaces, but at the cost of invasiveness and tissue damage that scale with the number of implanted electrodes^30,75–78^. Such systems are also difficult to remove or replace without causing damage to surrounding brain tissue. It is not yet clear whether approaches involving softer penetrating electrodes offer a substantially different tradeoff^30,75^. For this reason, non-penetrating cortical surface microelectrodes represent a potentially attractive alternative^69,79,80^. In practice, electrocorticography (ECoG) has already facilitated capture of high quality signals for effective use in brain–computer interfaces in several applications, including motor and speech neural prostheses^7,19,32,34,72,81–87^. Higher-spatial-resolution *micro*-electrocorticography (µECoG) therefore represents a promising combination of minimal invasiveness and improved signal quality. However, the limits of information that can be extracted at high-resolution have not been fully characterized.

Here we demonstrate a modular and highly scalable system of conformable, thin-film microelectrodes designed for rapid, minimally invasive deployment on the cortical surface. We demonstrate the practicality and *in vivo* performance of the system in Göttingen minipigs in anesthetized states and during awake locomotor behavior; as well as in the context of clinical neurosurgery for high-resolution cortical mapping during epilepsy surgery or to the removal of tumors near eloquent regions of the human brain associated with language and sensorimotor function. We describe implantation of thousands of electrodes simultaneously in multiple regions of the neocortex in both hemispheres, including areas related to vision as well as limb and facial somatosensory and motor function. We further demonstrate the use of these arrays for electrophysiologic functions required of contemporary brain–computer interfaces, including neural recording, cortical stimulation (“neuromodulation”), and neural decoding. We characterize the spatial scales over which electrophysiologic information is represented at the cortical surface, and show (contrary to some prevailing beliefs) not only that accurate neural decoding can be achieved, but also that accuracy improves as a function of both area coverage and spatial density. The platform is intended to facilitate human clinical use of brain–computer interface technology by delivering the microelectrode numbers and spatial densities required for advanced, high-performance brain–computer interface applications in a safe and time-efficient manner that is compatible with proven and reliable neurosurgical techniques.

## 2 Results

### System Overview

The system as configured for *in vivo* neural recording and stimulation comprises a modular set of thin-film microelectrode arrays designed for rapid, minimally invasive subdural implantation using a novel “cranial micro-slit” technique. Two versions of the microelectrode array were fabricated for this study. The first comprises 529 electrodes of multiple sizes ranging in diameter from 20 to 200 µm, while the second version comprises 1024 channels of three different diameters (977 at 50 µm and 42 at 380 µm, with an additional 5 reference electrodes at 500 µm). Microelectrode arrays can be inserted individually or in modular assemblies (see Supplementary Figure 1), with each array connected to a customized hardware interface. After subdural array implantation, the interconnecting cable of each microelectrode array module passes through a dural incision and a cranial micro-slit incision, is tunneled under the scalp as needed, and is connected to an individual head stage. The head stage contains electronics for analog-to-digital conversion and signal conditioning, and streams data to a custom software platform for real-time data visualization, processing, and storage. The overall system configuration is illustrated in Figure 1.

**Figure 1:**
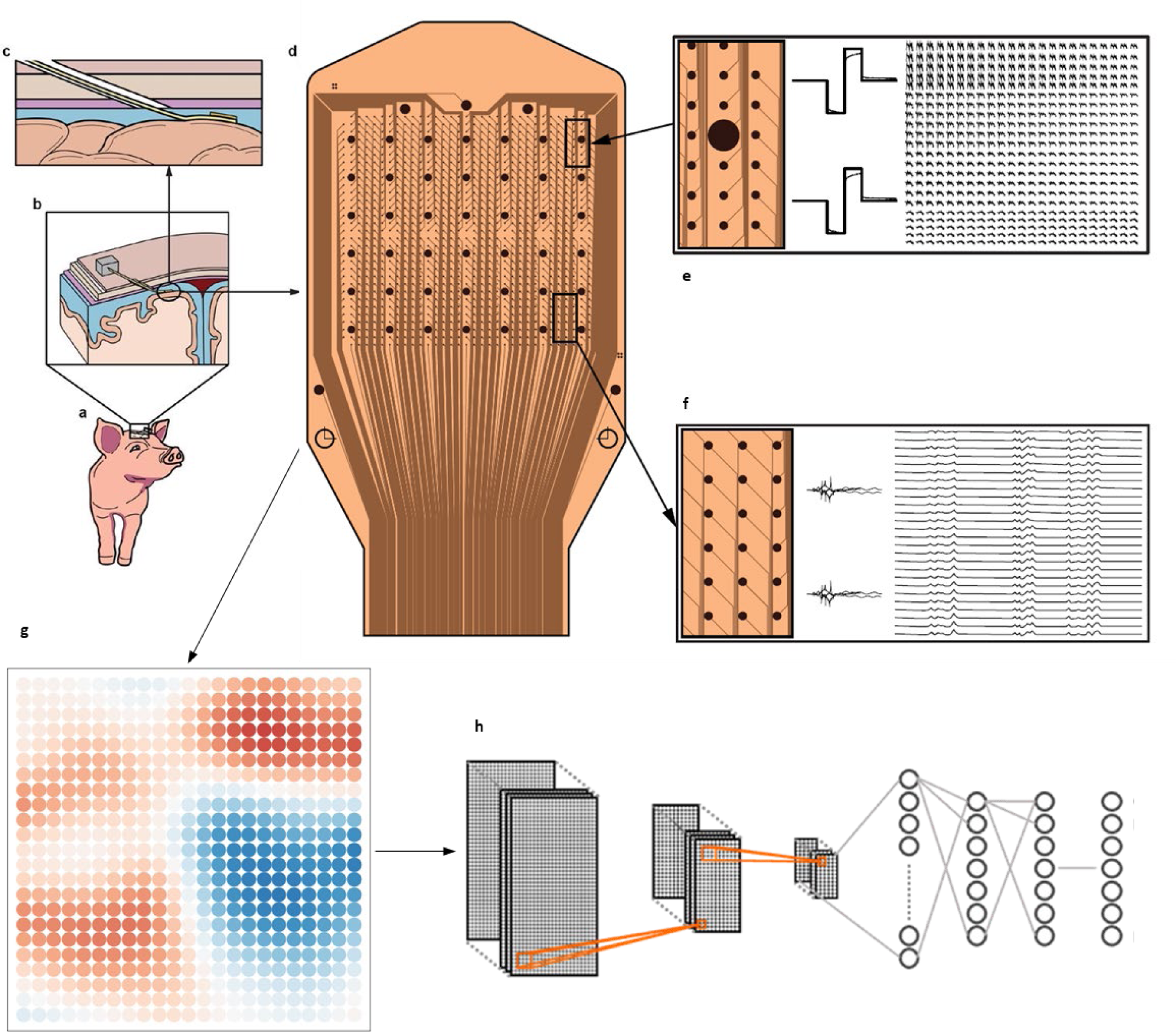
Overview of the minimally invasive thin-film neural interface system. This schematic illustration shows how neural activity is acquired from and stimulated by the thin-film cortical interface. (a) A Göttingen minipig undergoes cranial micro-slit implantation (b and c) of a set of subdural micro-electrocorticography arrays comprising a total of >2000 microelectrodes, in modules containing 529 (Supplementary Figure 2a) or (d) 1024 channels (see also Supplementary Figure 1) each. In the anatomic schematics (b and c) the subdural space is shown in blue, dura in purple, skull in beige. The outermost layer shown represents the scalp. (e) A representative 380 μm electrode is shown schematically together with example stimulation waveforms. (f) A group of 50 μm microelectrodes is shown in detail together with example traces from recorded neural activity. The microelectrode arrays provide (g) multichannel data that is used in a variety of electrophysiologic paradigms to perform neural recording of both spontaneous and stimulus-evoked neural activity as well as (h) decoding and focal stimulation of neural activity across a variety of functional brain regions.

### Electrode Array Characterization

All microelectrode arrays were thoroughly characterized prior to insertion testing through direct inspection and both *in vitro* and *in vivo* electrode impedance mapping. Process yield was >93% and 91% for the 529-channel and 1024-channel arrays, respectively. Electrode impedance exhibits a predictable dependence on electrode surface area, ranging from an average of 802 ± 30 kΩ for 20 µm electrodes to 8.25 ± 0.65 kΩ for 380 µm electrodes, and is robust to implantation, confirmed by the ratio of impedance before and after implantation showing little change across the array (Supplementary Figure 2).

### Minimal Invasiveness and Speed

We have demonstrated the safety and feasibility of inserting our high-density microelectrode arrays using a novel, minimally invasive “cranial micro-slit” technique (Figure 2, Supplementary Video 1). The procedure employs precision sagittal saw blades to make 500-to-900 micron-wide incisions in the skull at approach angles approximately tangential to the cortical surface, facilitating subdural insertion of our thin-film arrays without requiring a burr hole or craniotomy. Trajectory planning and insertion were performed using fluoroscopy or computed tomographic image guidance, and electrode insertion was monitored using neuroendoscopy. To validate the procedure, we have performed 22 cranial micro-slit insertions (between 1 and 4 insertions per animal) in 8 Göttingen minipigs (an additional 61 arrays were implanted through small craniotomies in 21 Göttingen minipigs for electrophysiologic and biocompatibility studies). The insertion time following cranial and dural incisions in human cadavers was consistently <20 s. For a single array of 1024 microelectrodes, this corresponds to an average insertion rate of <20 ms per electrode over 1.56 cm^2^ of cortical surface area. In addition, we have performed multiple cranial micro-slit insertions in 23 fresh cadaveric human heads, targeting the precentral gyrus in the expected region of upper extremity primary motor cortex, and verifying placement with a combination of endoscopy, fluoroscopy, and computed tomography (CT). We have demonstrated that the entire surgical procedure for cranial microslit insertion, from initial skin incision to endoscope-guided array placement and final securing of the array positions, can be safely performed in under 20 minutes.

**Figure 2:**
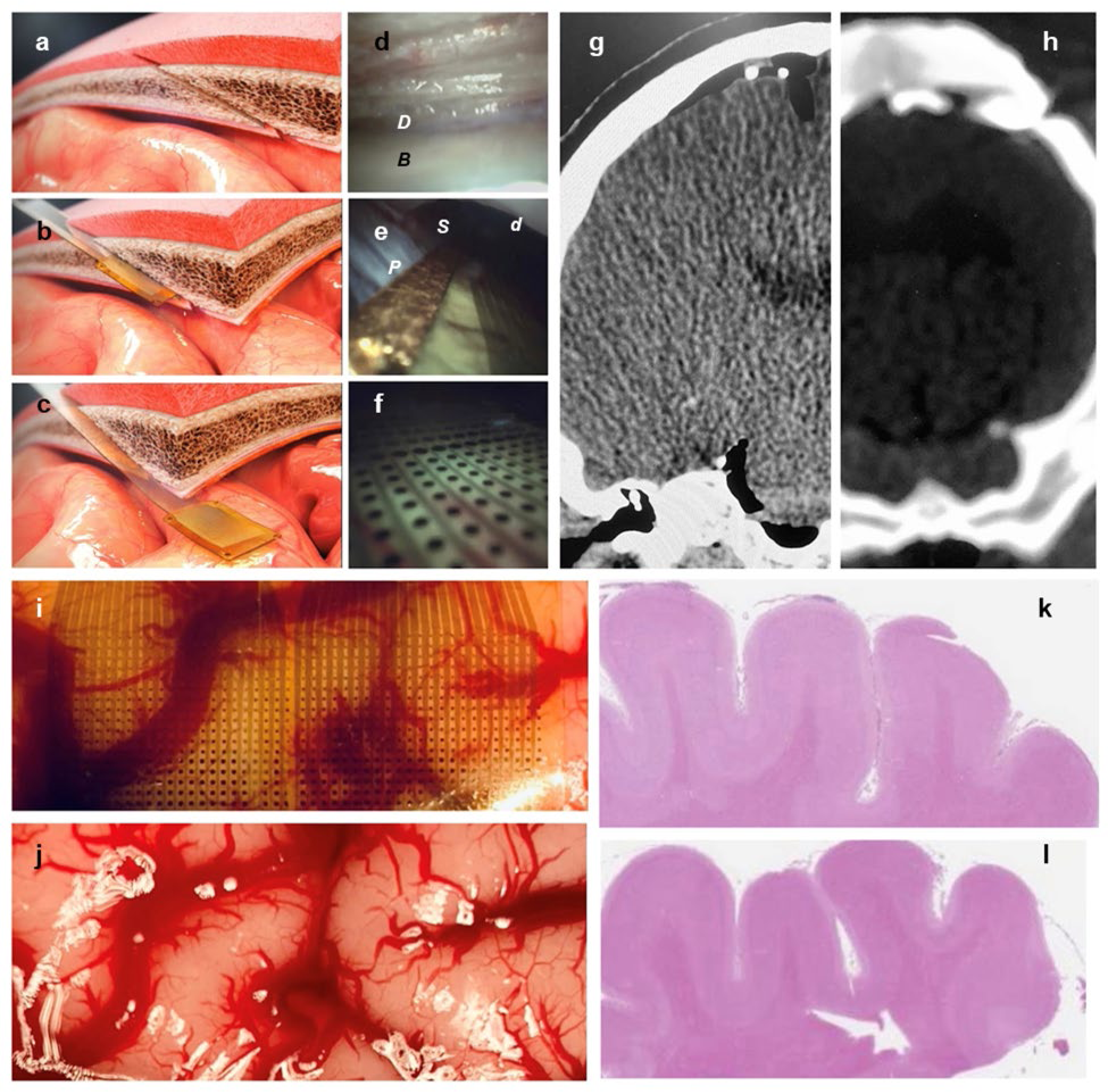
Safety and reversibility of neural interface implantation. The “cranial micro-slit” insertion technique is illustrated (a-c), showing guidance of the thin-film-based array into the subdural space. A scalp incision is made, followed by a cranial incision at an angle tangent to the cortical surface in the location targeted for array placement; the dura is then coagulated and incised (not shown). The trajectory of these aligned incisions is shown (a), together with an electrode array mounted on a stiffening stylet for subdural implantation through the cranial slit (b). The stylet inserts into a polyimide pocket on the back of the array. The array-stylet assembly is inserted into the subdural space. The stylet is then removed, leaving the array in place (c). Endoscopic views of the corresponding surgical steps are shown as performed in a human cadaveric head (d-f), showing the dura as observed through the microslit osteotomy (d), the array with its radiopaque gold marker being inserted into the subdural space (e), and the microelectrodes *in situ* with the cortical surface seen through the thin film array (f) (*B*, cut surface of skull seen from within the cranial micro-slit; *D*, outer surface of the dura; *d*, undersurface of the dura; *S*, subdural space; *P*, pia of the cortical surface). (g-h) Computed tomography (CT) scans in the coronal plane are shown of arrays placed via cranial micro-slit technique on human primary motor cortex (g) and pig somatosensory cortex (h), with array edges apparent due to the presence of radiopaque gold markers. (i-j) A modular configuration of two 529-channel arrays (1058 microelectrodes in total) is shown *in situ* on the cortical surface following a small frontal craniotomy in a Göttingen minipig (i); the same region of cortical surface is shown immediately following array removal (j), demonstrating an intact pial layer and no damage to the brain. (k-l) Safety was also assessed using standard, semi-quantitative histology techniques following 30-day chronic array implantation. Histologic analysis demonstrated microscopic preservation of the cortical surface architecture and no systematic differences between implanted and non-implanted control regions (k and l, respectively, hematoxylin and eosin).

### Safety and Reversibility

We assessed the safety and reversibility of microelectrode array placement by gross and microscopic examination of cortical insertion sites following implantation (Figure 2). Insertion sites were inspected by craniectomies immediately postmortem, following euthanasia, to assess the cortical surface for evidence of tissue damage following array removal (“reversal” of implantation). An example image of the cortical surface after array removal and postmortem craniectomy is shown in Figure 2j. Conformal electrode array contact caused no tissue disruption at the cortical surface. Histologic evaluation of cortical sections immediately beneath implantation sites following acute (≤8 h) and chronic (>30 day) implants revealed no evidence of acute or chronic tissue injury (see Figure 2k-l).

### Modularity and Scalability

The fabricated arrays facilitate alignment and modular assembly, as shown in Supplementary Figure 1, so that multiple electrode modules can be joined to yield larger constructs covering larger portions of the cortical surface, without significantly increasing the complexity, risk, or time required for array insertion. It is also possible to insert multiple arrays through a single slit. We performed *in vivo* insertions of doubly connected 529-channel modules (1,058 channels over 0.96 cm^2^ of cortical surface area).

We have also demonstrated the ability to interface with multiple anatomic and functional areas of the neocortex simultaneously *in vivo*. We have performed simultaneous bilateral insertions over somatosensory and motor cortex, and have recorded simultaneously from multiple sensory regions including portions of somatosensory, visual, and auditory cortex (Supplementary Figure 3). Functional localization for each region is confirmed using evoked potentials (Figure 3).

**Figure 3:**
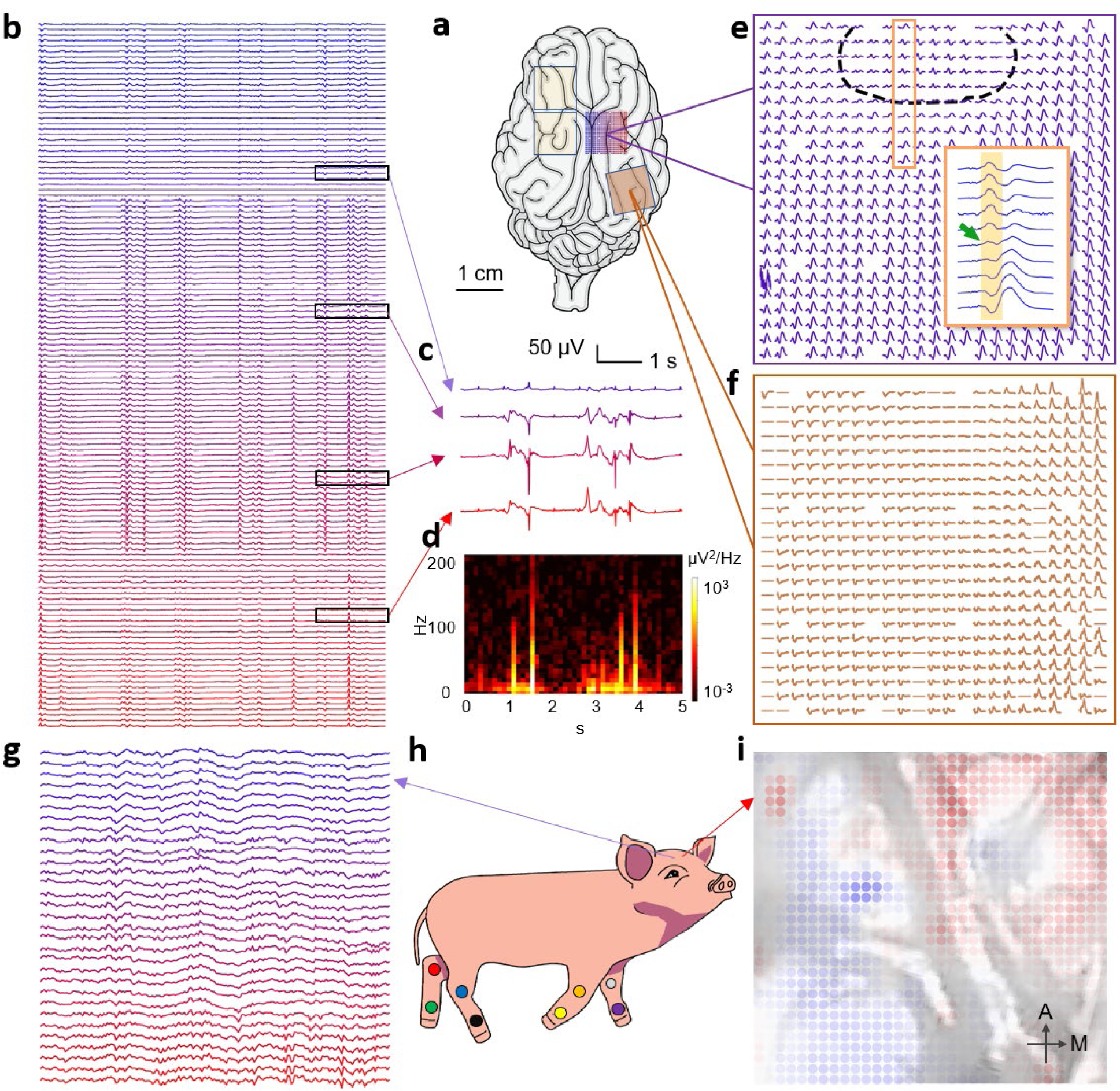
Neural recordings from multiple functional brain regions resolve electrographic features from the cortical surface at high spatial resolution in both anesthetized and awake-ambulatory states. (a) Schematic of the Göttingen minipig brain showing color-coded areas corresponding to placement of thin-film subdural microelectrode arrays. (b) Recording of spontaneous neural activity from somatosensory cortex in a sample of channels from the 529-channel array highlighted in (a). (c) Example recording traces, and (d) example spectrogram corresponding to the highlighted channels from an array over the right somatosensory cortex. (e) Somatosensory evoked potentials corresponding to electrical stimulation of the tibial nerve. The inset highlights reversal of phase over a sub-millimeter scale, and the ellipsoid highlights the two-dimensional nature of the isoelectric contour. Traces span 1 s and the maximum peak-to-peak signal is approximately 80 μV. (f) Visual evoked potentials from photostimulation of the left eye. (g) Example voltage traces obtained during spontaneous walking from one of two 1024-channel thin-film subdural microelectrode arrays placed over the sensorimotor cortex (one over each hemisphere) in a Göttingen minipig. (h) Schematic representation of the experimental setup, in which an adult Göttingen minipig was permitted to walk ad lib on a treadmill shortly after emerging from anesthesia following array placement. Accelerometers and motion-capture fiducials were placed on each limb in order to facilitate motion tracking time-synchronized with cortical activity. (i) Representation of cortical surface activity from the left hemisphere sensorimotor cortex, as recorded from a 1024-channel electrode array, displayed as a color-map of voltage, superimposed on and aligned to an image of the underlying cortical surface (each dot corresponds to one microelectrode in the array).

### Neural Recording

We sought to leverage the high resolution and data bandwidth of our integrated system to map electrocortical activity on a fine-grained scale (Figure 3). In preclinical studies involving Göttingen minipigs, implanted arrays were used for high-bandwidth and high-spatial-resolution neural recording of both spontaneous cortical electrographic activity and evoked potentials from multiple functional regions (Figure 3). During free recording of spontaneous cortical activity, our software enables real-time visualization of raw voltages or spectral power of 1,024 channels simultaneously (see Figure 3, Supplementary Video 2, Supplementary Video 3, and Supplementary Video 4). Recorded electrocortical activity from individual channels can be viewed in both time and frequency domains, revealing the presence of electrocortical activity at frequencies up to 500 Hz. The degree of correlation across electrodes decreases with distance and with increasing frequency (Supplementary Figure 4). Importantly, even at 300 μm spacing, adjacent electrodes exhibit incompletely correlated activity, particularly at higher frequencies. For example, beta-band *r*^2^ is in the range of 0.45±0.03 at 300 μm spacing for 50 μm electrodes, suggesting that even at this spatial scale the total amount of electrophysiologic information available at the cortical surface has not been completely extracted.

To further explore the utility of high-spatial-density neural recording, evoked potentials were obtained across multiple arrays and multiple functional regions. Robust somatosensory evoked potentials (SSEPs) were obtained in arrays positioned over the somatosensory cortex following electrical or tactile stimulation (Figure 3) of all 4 limbs. When the arrays span both motor and sensory cortex, the SSEPs demonstrate clear phase reversal at the motor-sensory junction (Figure 3e, inset); in contrast to traditional macroelectrode strips, which enable only coarse localization of the boundary to within a fraction of a centimeter, and typically in one dimension, we are able to identify this boundary with 300 μm resolution, and as an isoelectric contour line in two dimensions, providing precise mapping of functional boundaries on the cortical surface. Robust visual evoked potentials (VEPs) were similarly obtained in arrays positioned over the visual cortex following time-synchronized photostimulation of the retina (Figure 3f). We also recorded electrocorticographic activity in awake, freely ambulating animals (Figure 3g-h). Time-synchronized neural data from 2048 channels (1024 per hemisphere in the region of primary sensorimotor cortex) was acquired together with three-dimensional motion capture data using multiple fiducial markers on each limb (Figure 3h) as well as accelerometers mounted on all four limbs and the head.

### Multi-Modal Neural Decoding

We next sought to evaluate the ability of the system to use high-resolution electrocortical activity to perform neural decoding in a variety of paradigms related to somatic sensation (multi-point discrimination), vision, and volitional walking during conscious awake behavior (Figure 4).

**Figure 4:**
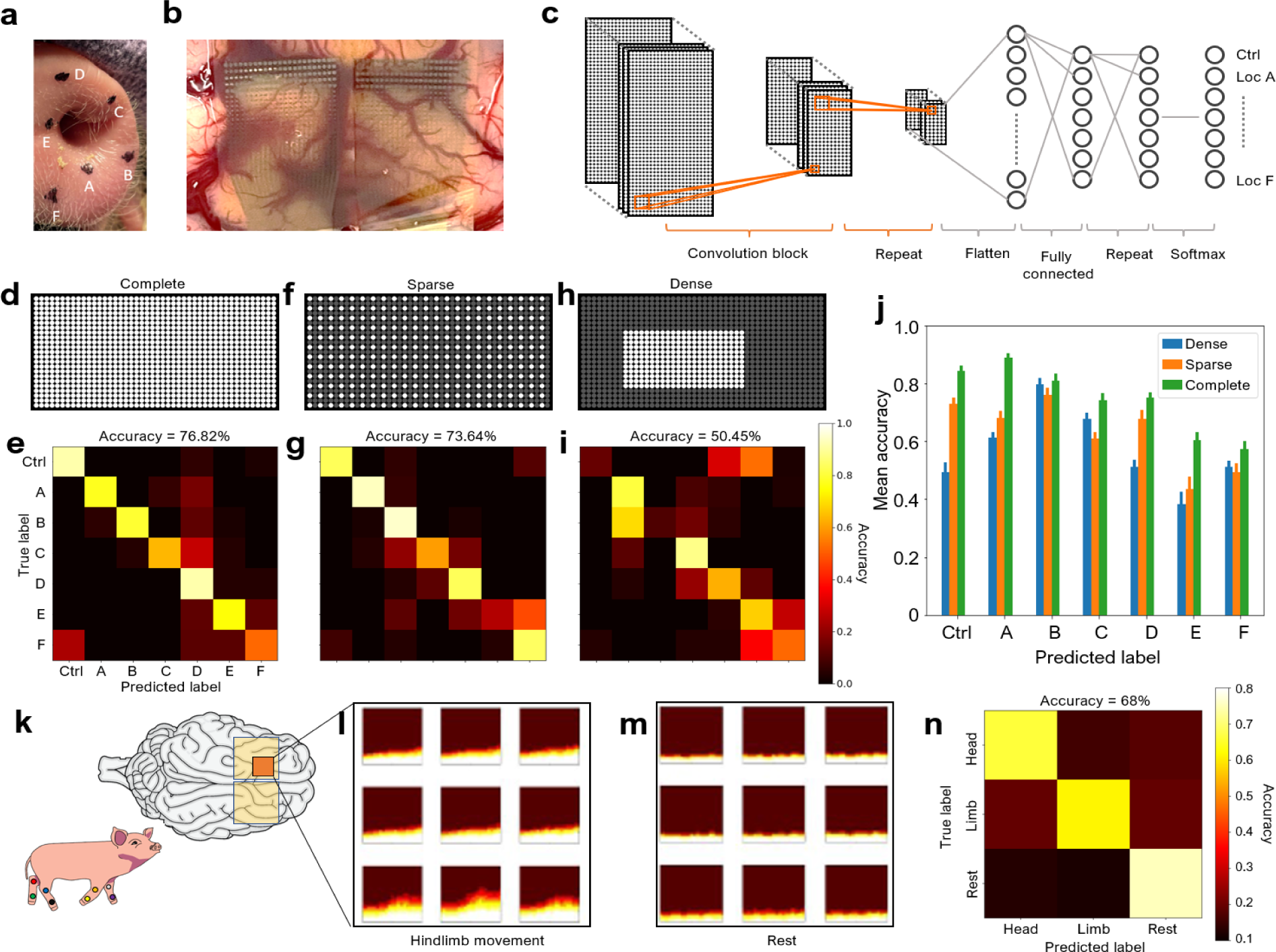
Neural decoding accuracy benefits from both greater extent of cortical surface area and higher density of microelectrode coverage. (a) Tactile stimulation locations on the rostrum of the Göttingen minipig used for neural decoding. (b) Placement of two electrode arrays (1058 microelectrodes total) on the cortical surface overlying the rostrum somatosensory cortex. (c) Baseline architecture of the convolutional neural network used for decoding stimulated location. A convolution block consists of the following steps in sequence: 3×3 convolutions, leaky rectified linear unit (ReLU), dropout during training, “max pooling” with stride 2, and batch normalization. Each fully connected layer is followed by leaky ReLU. (d) All electrodes are recruited in decoding (“complete grid”). (e) Confusion matrix of decoding with the entire array. (f) Every other electrode in both spatial dimensions is recruited for decoding (“sparse grid”), totaling 23×12 electrodes. (g) Confusion matrix of decoding with sparse grid. (h) A 23×12 grid of adjacent electrodes (“dense grid”) is recruited for decoding, resulting in a data rate equivalent to using a sparse grid. (i) Confusion matrix of decoding with dense grid. (j) Decoding accuracies for each stimulation location and control, using convolution kernels emphasizing different options provided by the high-density cortical surface arrays (complete coverage at full density, sparse density, and subsampled partial coverage). Error bars represent the standard error of the mean for 25 models trained per grid configuration. (k) Schematic showing bilateral placement of 1024-channel electrode arrays (2048 microelectrodes total) for motor decoding during volitional walking (inset schematic). (l and m) Illustrative subsamples of spectrograms drawn from the same group of electrodes across the array (highlighted orange region in the schematic of (k)), corresponding to each of two forelimb movement states, moving (l) and stationary (m), demonstrating distinct, state-dependent patterns of cortical activity. (n) Decoding of limb movements from pre-movement microelectrocortical activity. Confusion matrix of decoding limb movement and rest state using pre-movement neural activity with a convolutional neural network (CNN); 10-fold cross-validation.

In somatosensory decoding experiments designed to assess multi-point discrimination, two 529-channel arrays or one 1024-channel array was placed over the somatosensory cortex and electrocortical activity was recorded during manual stimulation of predefined rostrum locations separated by <1 cm (Figure 4a-b). The observed evoked potentials from different stimulation locations exhibited spatial and temporal variation on multiple scales (Supplementary Figure 4 and Supplementary Figure 5). Multi-scale spatial variation was also observed in free recording, especially when comparing across EEG bands (Supplementary Figure 4). We therefore developed a custom deep neural network architecture for decoding that incorporates both temporal and spatial features, consisting of multiple convolutional layers and several fully connected layers (Figure 4c). The network takes as input band-passed filtered voltages from 1058 spatially distributed electrodes over 200 consecutive time samples (representing 200 μs at 1 kHz sampling) and predicts one of 7 stimulation classes (one for each somatosensory stimulation site plus a control class corresponding to absence of tactile stimulation). Across multiple independent animals, we trained a custom network on 80% of the data and evaluated the accuracy of the resulting model on the remaining 20% of the data. We obtained overall test-set accuracies of 66–76%, with most of the errors being near-diagonal, reflecting adjacent-position mismatches (Figure 4 and Supplementary Figure 6). By comparison, using an identical architecture on coarse decoding tasks resulted in accuracies approaching 100% for full-field visual decoding and 80–85% for limb SSEP (Supplementary Figure 7).

We next investigated the relative importance of spatial coverage and channel resolution by down-sampling the rostrum stimulation electrode array data in two different ways (Figure 4d-j): in dense sampling, we trained a model using only one quarter of the electrodes in the region corresponding to maximum saliency, while in sparse sampling, we alternatively dropped every other row and column of the electrodes. For most rostrum stimulation positions, sparse sampling performed better than dense sampling, however, for two of the stimulation locations, dense sampling was superior. Utilizing the complete array outperformed either approach for all stimulation locations, highlighting the need for both broad spatial coverage as well as high resolution to obtain accurate neural decoding.

Inspired by these findings, we developed a new deep learning architecture (DenseSparseNet) that incorporates information from multiple spatial scales by performing convolutions on the full, half-, and quarter-down-sampled arrays; this architecture improved decoding accuracy by approximately 10% in all three experiments, resulting in accuracies across experiments ranging from 77–86% (Figure 4e), and further highlighting the advantages of a scalable, high-resolution surface array for high sensitivity neural decoding.

We also sought to decode volitional motor activity in awake, consciously behaving animals. To this end, we performed bilateral craniotomies and placed one 1024-channel array over each motor cortex (for a total of 2048 electrodes per animal) and awakened the animal in a harness suspended over a treadmill designed to permit walking *ad lib* (see Methods). The spontaneous motor activity of the animal was recorded using motion-tracking cameras and a 3-axis accelerometer on each limb. We observed characteristic and reproducible electrocortical spectral changes in periods immediately preceding voluntary limb or head movement, distinguishable from rest (Figure 4k-n). We were able to resolve gross features of motor behavior consistently across different animals, demonstrating three-class decoding for head, limb movement, and rest with a simple CNN architecture (see Methods) obtaining accuracies of 53–69% (*k*-fold cross validation; *k*=5, 10). Furthermore, depending on the behaviors exhibited by each animal, we also applied some individualized behavior decoding paradigms, showing that versatile decoding of sensory and motor modalities from multiple body regions could be achieved with the same array and network structure (Supplementary Figure 5) (60-80%, *k*-fold cross validation; *k* =5).

In a pilot clinical study involving five neurosurgical patients undergoing intraoperative electrophysiologic mapping, we further evaluated the ability of our integrated system to acquire, process, and display high-spatiotemporal-resolution electrocortical data in real time. Spontaneous electrocortical recordings were obtained in two patients under general anesthesia, on conventional 4-electrode strips and in higher resolution on the 1024-electrode devices. Where a reversal of phase in the SSEP, corresponding to the functional central sulcus, was demonstrated using the conventional 4-electrode strip, this was recapitulated in higher resolution using the 1024-electrode device, revealing a high-resolution phase reversal “contour line” in two dimensions over the cortical surface. Three patients underwent awake language mapping, and 1024-channel electrocortical recordings were obtained from the left superior temporal gyrus, left angular gyrus, and left frontal operculum, respectively, during speech time-synchronized to auditory or single-word visual cues. We were able to decode speech events from the electrocortical activity of these awake neurosurgical patients (Figure 5a-h). As expected in the context of intraoperative language mapping in awake patients, the spectral properties of the electrophysiologic and speech signals were strongly correlated. Compared to intervals without speech attempts, we observed reproducible modulation of low-frequency (0–40 Hz) bands in the intervals immediately surrounding actual speech production events (Figure 5c-g, Supplementary Video 4). These strong correlations provided a basis for binary neural decoding of speech onset, achieving an accuracy of 79.8% (95% CI: 77.6–82.0%) from even the short training windows provided during awake craniotomies. Using only 4 minutes of training data containing 54 patient utterances, we demonstrate 79% offline decoding accuracy in distinguishing speech events from silence on the basis of the electrocortical activity from the high-resolution array (Figure 5h).

**Figure 5.**
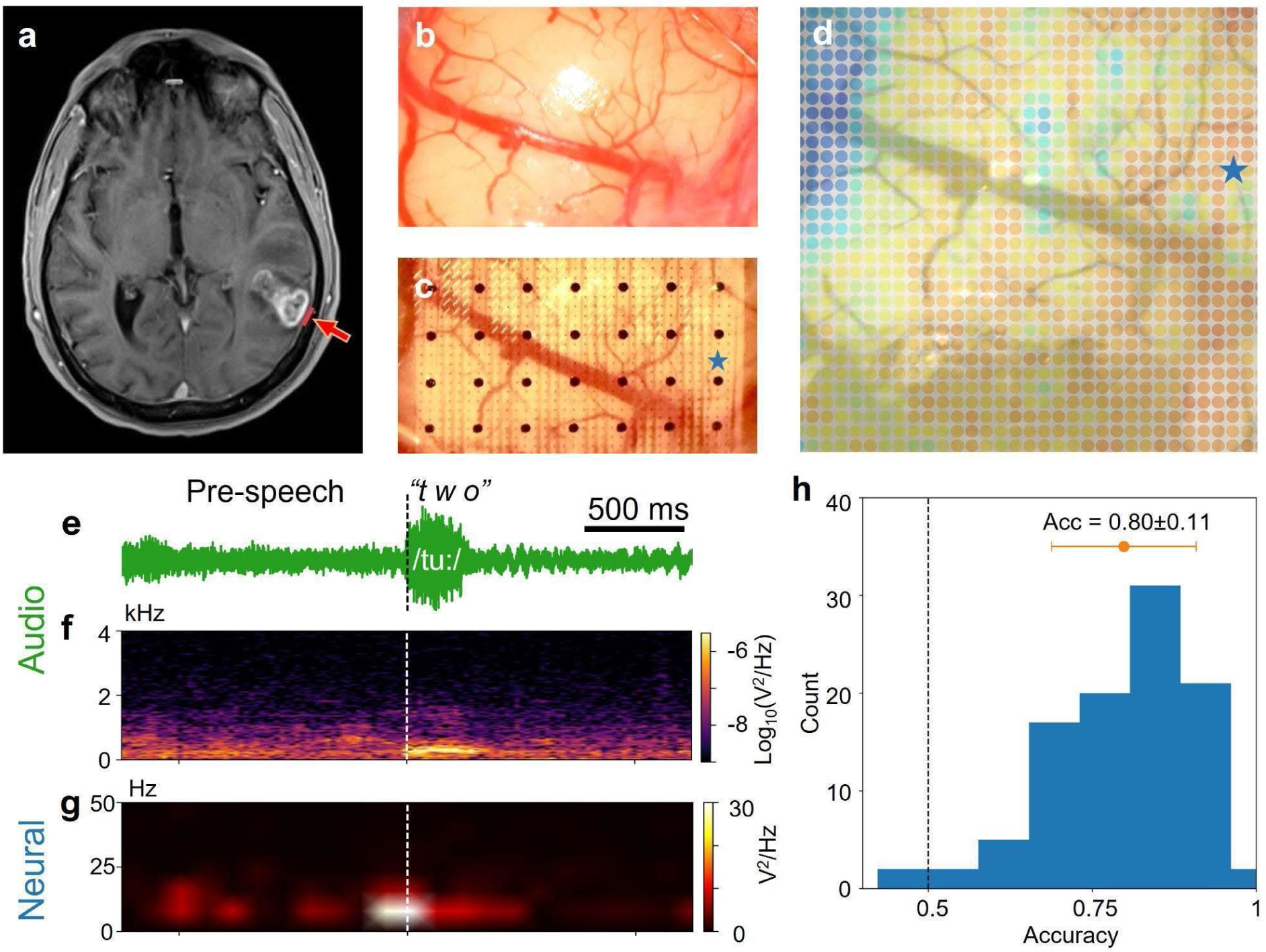
Human intraoperative electrocorticography. (a) Axial gadolinium-contrast-enhanced T1-weighted MRI of the brain of one patient involved in the pilot study, demonstrating a tumor in the left temporal lobe. The red bar indicates placement of a 1024-channel electrode array during awake language mapping. Cortical surface of the left superior temporal gyrus prior to (b) and following (c) electrode array placement. (d) Overlay-representation of electrocortical activity from the 1024-channel electrode array at the time point indicated by the dashed line in (e-f), immediately prior to speech onset. The color map represents normalized raw voltage as obtained from the digital steps of the analog-to-digital converter (*dark blue*, low; *yellow*, high). (e) Audio amplitude recorded during patient speech. (f) Time-frequency spectrogram of audio recording during the same time interval shown in (e). (g) Time-frequency spectrogram of the voltage waveform from a representative electrode *(star)*. (h) Accuracy of decoding speech onset on the basis of a 4-minute training set of spoken words, in offline decoding, as assessed in 100 randomly shuffled train-test samples.

### Neural Stimulation

The electrode arrays are capable of bidirectional function, with every electrode able to perform either recording or stimulation. To characterize the ability of our system to modulate cortical activity *in vivo*, 16 electrodes per 529-channel array were designated for use in cortical stimulation. Safe stimulation thresholds were determined *in vitro* (50–100 µA, 200 µs pulse width; see Methods) and cortical stimulation using these parameters was performed *in vivo*. The 200 µm electrodes were used for stimulation in each trial, and the remaining sites on the same array, as well as all sites on adjacent arrays, were used for recording.

We performed focal stimulation of the visual cortex using the paradigm described, while recording stimulus-evoked cortical activity (Figure 6). Cortical stimulation modulated high-gamma-band power in a focal region surrounding the stimulated electrode over timescales of >2 s (Figure 6e, g, h, and i). Importantly, this effect was reduced by increasing the depth of anesthesia (Figure 6f and j), and the effect of stimulation was observed to spread across array boundaries, indicating that the observed effects are physiological and not artifacts of stimulation. Within the range of parameters tested, no induced saccades were observed in these experiments.

**Figure 6:**
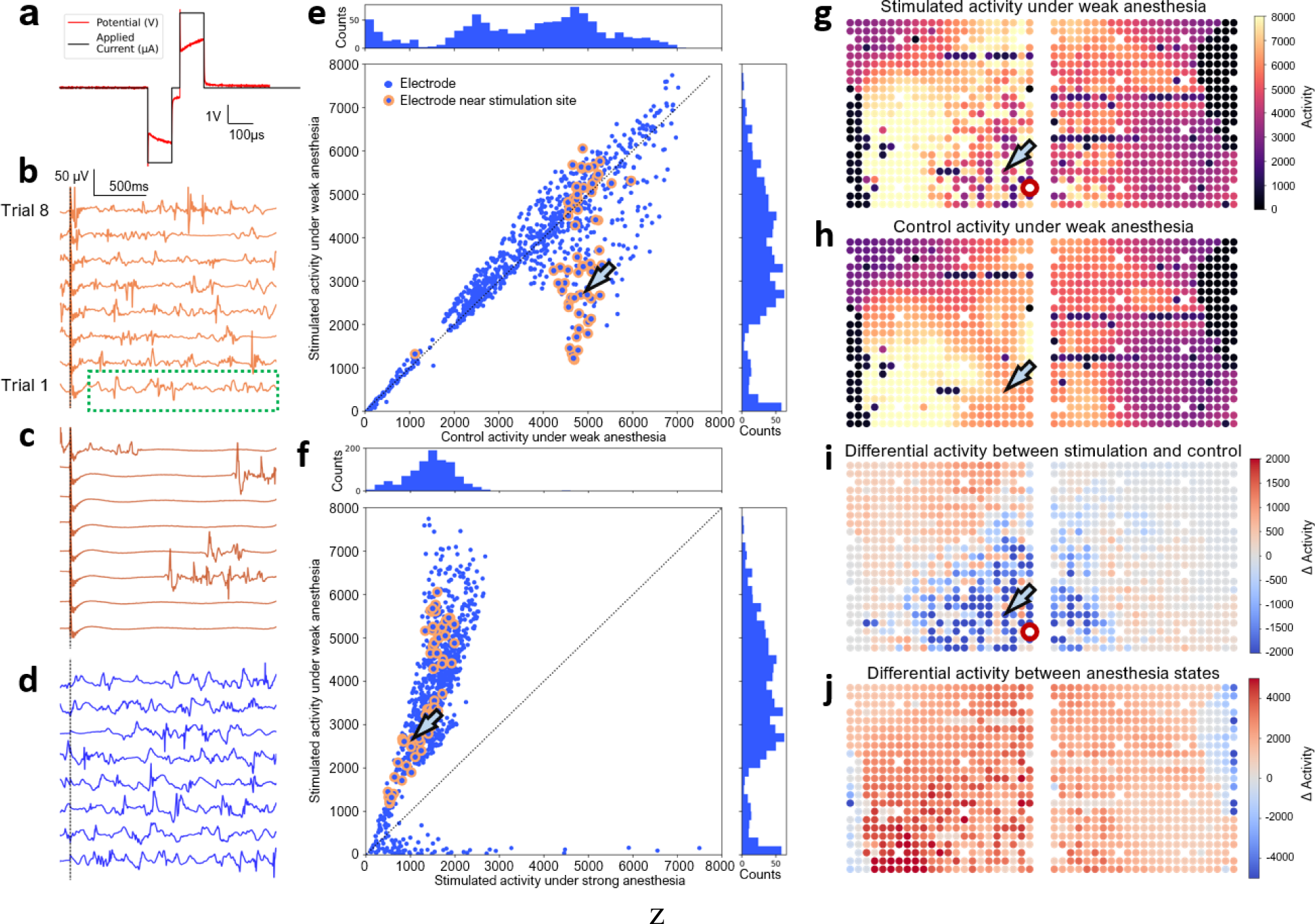
Cortical microstimulation modulates cortical activity in ways that can be characterized in high spatial and temporal detail. (a) Stimulation waveform used for *in vitro* confirmation of safe polarization potential, with 100 μA overlaid on the waveform for reference. *In vivo* applied current waveforms used the same applied current but without the interphase delay used for identifying polarization potential. (b) Example traces for an electrode (blue arrow in g) near the stimulation electrode (red ring in g) for 8 stimulation trial recordings. The “activity” of each trial is computed as the variance of the signal from 200 ms to 2000 ms post-stimulation (green box), and the average activity is taken over 40 trials. (c) Corresponding traces for the animal under heavier anesthesia. (d) Corresponding traces for the electrode without stimulation under light anesthesia. (e) Stimulated activity is plotted against control activity. Each point represents an individual microelectrode; the highlighted points are located within 5 electrode spacings from the stimulating electrode. The histograms show the distributions of activity with (side panel) and without (top panel) stimulation. Blue arrow shows the same electrode as indicated in panel (g). (f) Stimulated activity under light versus heavy anesthesia, plotted using a scheme analogous to that used for (e). The histograms show the distributions of stimulation-induced activity under different levels of anesthesia. (g) Activity across the two adjacently placed arrays with stimulation applied at the highlighted electrode. (h) Activity across the arrays without stimulation, using the same color scale as in (g). (i) Differential activity across the arrays, calculated as the difference between (g) and (h), revealing a region of suppressed activity surrounding the stimulating electrode. (j) Map of differential stimulated activity between light and heavy anesthesia.

## 3 Discussion

Here we have described a modular and highly scalable brain–computer interface platform capable of rapid, minimally invasive surgical deployment over multiple areas of the cortical surface in a reversible and atraumatic manner^88^. Through a series of live animal and human cadaveric surgeries, we demonstrate the safety and feasibility of delivering >2,000 microelectrodes into the subdural space of multiple functional areas of the brain at a rapid rate (>1,000 electrodes per minute) through 400-to 900-micron wide skull incisions. Using thin-film microelectrode arrays that achieve 200-to-1,000-fold higher electrode density than standard cortical grids, we definitively demonstrate that electrophysiologic information is available at the cortical surface at scales as small as <300 µm, and we show that both high density and wide spatial coverage are required to achieve accurate neural decoding from electrocortical activity. Finally, we demonstrate the ability of these bidirectional electrodes to modulate cortical activity through focal stimulation of individual electrodes. The platform is designed to deliver the large microelectrode numbers and high spatial densities required to facilitate confident adoption and routine clinical use of advanced, high-performance brain–computer interface technologies. Our early experience in human patients substantiates the clinical utility and usability of the system.

Clinically useful neural interfaces must balance the need for improved function, achieved through increasing channel counts and spatial resolution, against the invasiveness of the surgical procedure and damage to brain tissue associated with penetrating intracortical electrode arrays. In designing our system, we have chosen to prioritize safety and scalability over other design criteria with an eye toward rapid and efficient use in human clinical applications. While it is unlikely that any single system or electrode type will be ideally suited to the full breadth of future clinical applications of neural interfaces, safety, scalability, and reversibility are key features to maximize the addressable clinical populations that could benefit from this technology in the near future. Specifically, it is important that tissue damage and total implantation time for neural interfaces not increase in proportion to electrode count, as scaling such systems by several orders of magnitude would then lead to prohibitive levels of tissue damage^89–92^ or impractically long implantation times. The Layer 7 Cortical Interface system described here compares favorably on these measures, enabling thousands of electrodes to be deployed within minutes to multiple functional areas without damaging cortical tissue. Indeed, it becomes conceivable to envision deploying a thin-film-based neural interface over the majority of the accessible human neocortex.

Several groups have used advanced surface array techniques to correlate neural activity with motor function for the control of neural prostheses in paralyzed patients^7,84,85,93–96^, to achieve speech decoding in anarthric patients^6,19,80,86,97,98^, or for other applications of high-resolution electrocorticography^72,80,99–109^. Using the significantly increased resolution of the Layer 7 microelectrode array, we demonstrate neural decoding across multiple functional modalities, including gross and fine somatosensation, vision, and volitional motor function during awake, spontaneous, untrained behavior, as well as human speech, with a machine learning framework that suggests that work in the field to date has not yet fully capitalized on the electrical information present at the cortical surface. Maximal decoding performance is achieved by incorporating information at multiple spatial scales, requiring systems and techniques that can combine wide coverage and high spatial resolution. Additionally, we have demonstrated the feasibility of using these electrode arrays for dynamic, sub-millimeter-scale mapping of the cortical surface in clinical practice, in support of high-precision neurosurgery in eloquent brain regions.

Precision neuromodulation is also a key capability of closed-loop brain–computer interfaces, as well as future neural prostheses for restoring functions such as somatic sensation, proprioceptive feedback, and vision. Here we characterize a safe operating regime for cortical stimulation from thin-film cortical surface microelectrodes, stimulate the cortical surface, and monitor stimulus-evoked cortical activity in a manner that facilitates visualization and analysis of cortical surface activity at spatial and temporal resolutions and total spatial extents not previously possible.

The system we describe here, the Layer 7 Cortical Interface, is a modular, scalable, minimally invasive brain–computer interface platform. The system is designed to deliver the benefits of high-density, high-channel-count, high-data-rate neural interfaces to the millions of patients with neurologic disorders who stand to benefit from this technology.

## 4 Methods

### Array Fabrication and Characterization

Initial 529-channel microelectrode arrays were fabricated on 6–8” wafers using a spin-on polyimide. The fabrication process briefly comprised spin-coating, soft-bake, and vacuum cure of an approximately 10 μm layer of polyimide; photolithographic patterning, deposition, and liftoff of 20 nm/210 nm/20 nm Ti/Pt/Ti trace metal; O_2_ plasma treatment of the polyimide surface; spin-coating, soft-bake, and vacuum cure of an approximately 10 μm layer of polyimide; hard mask deposition and patterning for polyimide outline and electrode site opening; polyimide etch and electrode surface exposure in O_2_/CF_4_ plasma; hard mask strip; photolithographic patterning, deposition, and liftoff of 20 nm/20 nm/500 nm of Ti/Pt/Au bond pad metallization; and O_2_ plasma post-clean of the polyimide surface. Following microfabrication, devices were released in deionized water, optically inspected for trace, electrode, and pad defects, dehydration baked, and thermocompression bonded to an organic interposer using a flip-chip tool. The fabrication process is similar for the 1024 channel array, except that the first metal stack is adjusted to include gold in the trace metal stack for reduction of trace impedance (with platinum remaining as the tissue contacting material), and designs are adjusted as pictured in Figure 1 and Supplementary Figure 1. Key design changes include 50 μm and 380 μm recording and stimulation electrodes, respectively, as well as 500 μm on-array reference electrodes.

Additional microfabrication details have been described in our prior work^71^.

Each array pocket was laser cut from adhesive-backed polyimide film, then aligned with the 800 µm alignment holes and markers at the distal tip of the microelectrode arrays and compressed to form the final structure used for insertion.

The microelectrode arrays are designed to be assembled into larger connected modules in a scalable fashion to achieve greater cortical coverage. Spacing and orientation were controlled during modular assembly with the assistance of alignment holes. The arrays were bonded by applying ISO 10993 biologically tested ultraviolet-curing cyanoacrylate to the overlapping regions of adjacent array modules.

Prior to assembly, bonded microelectrode array-interposer assemblies were optically inspected in bond, cable, and electrode areas, and a sampling of electrodes were characterized electrochemically. Electrochemical characterization was performed on a potentiostat (Wavedriver 100, Pine Research, Durham, North Carolina, United States of America) in a 3-electrode configuration (with Ag/AgCl reference electrode and Pt coil counter-electrode), and comprised cyclic voltammetry (CV) and electrochemical impedance spectroscopy (EIS) on at least one electrode per size in phosphate buffered saline (PBS) at pH 7.4. The CV measurements were performed (from 0 to 1.2 V to -0.65 V to 0 V relative to the reference electrode) to confirm electrode surface identity using platinum oxidation and Pt-O reduction peaks, hydrogen adsorption, and H_2_ oxidative desorption. In addition, CV measurements provide information on charge storage capacity and real surface area and identify the water window. EIS measurements were performed from 10 Hz to 10 kHz (on each electrode size) to confirm that 1 kHz impedance and cutoff frequency are within expected ranges, and to provide references for later *in vitro* impedance mapping performed using the Intan chips in a two-electrode configuration. *In vitro* impedance mapping was performed in PBS on fully assembled devices (across all electrodes) at 100 Hz, 200 Hz, 500 Hz, 1,000 Hz, 2000 Hz, and 5,000 Hz using the Intan chips in our custom 529- and 1024-channel head stages.

### Surgical Implantation

#### Minimally Invasive Surgical Implantation

*In vivo* testing of the minimally invasive surgical insertion technique and electrode array performance were performed in adult female Göttingen minipigs. The breed was selected for well characterized functional neuroanatomy as well as skull thickness comparable to that of adult humans. The study protocol was approved by the IACUC of DaVinci Biomedical Research Products, Inc. Local anesthesia was achieved in the region of the skin incisions using intradermal lidocaine. General anesthesia was maintained with isoflurane at levels sufficient to produce analgesia without suppressing electrocorticographic activity, a balance that was facilitated by the minimally invasive nature of the procedure.

We developed a “cranial micro-slit” technique for array implantation. In order to insert each electrode array, a cranial incision was made using a modified 400-micron thick sagittal saw blade (or an 800-micron thick pair of such blades), at an entry angle tangential to the cortical surface. A 350-micron fiber-scope was then inserted through the cranial incision and used to visualize the dura, which was coagulated and cut under direct endoscopic vision. Endoscopy was similarly used to guide insertion of each electrode array into the subdural space. In some instances, a 1.6 mm semi-rigid endoscope was used through a separate pilot hole to facilitate improved image quality for photography or videography of the procedure.

Electrode arrays were positioned subdurally on the cortical surface under simultaneous endoscopic and fluoroscopic guidance. Manipulation of each thin-film array was performed using a radiopaque stylet. The stylet tip was designed to fit within a polyimide “pocket” on the reverse side of each array. Placement, depth, and angulation of cranial incisions and electrode arrays were also guided by fluoroscopy or computed tomography (CT). Each stylet was removed following fluoroscopic confirmation of array position, leaving only the thin-film subdural microelectrode arrays in position on the cortical surface.

In order to decouple assessment of the minimally invasive surgical technique from characterization of surface microelectrode array recordings, additional procedures were performed in which electrode arrays were placed on the cortical surface through small, traditional craniectomies. In these procedures the craniectomy was performed with a high-speed burr, the dura was separately incised and elevated to expose the cortical surface in the region of interest, hemostasis was meticulously achieved, and the microelectrode was placed on the cortical surface under direct vision.

#### Human Intraoperative Array Implantation

On the basis of the reversibility of the electrode array deployment and existing safety and biocompatibility data, the Layer 7 Cortical Interface was designated a “Non-Significant Risk” device in the context of limited intraoperative use, and Institutional Review Board (IRB) approval was obtained for short-duration cortical surface recordings alongside standard electrophysiologic mapping performed according to the neurosurgical standard of care, with informed consent obtained preoperatively (West Virginia University Medical Center IRB Protocol Number 2207618749). The human electrophysiologic data reported here were obtained under total intravenous anesthesia with propofol and fentanyl or propofol and remifentanil, with the addition of dexmedetomidine in patients undergoing awake language mapping, with a 1024-channel microelectrode array placed alongside a standard subdural electrode strip for up to fifteen minutes. In these patients the subdural electrodes were placed after traditional craniotomies were performed to expose the regions of surgical interest.

### Electrophysiology

#### System Configuration and Recording Hardware

The 529-channel customized neural recording and stimulation system is based on chips and controllers made by Intan Technologies (Los Angeles, California, United States of America). The custom amplifier printed circuit boards (PCBs) used to interface with the implanted electrode arrays each contained eight of the RHD2164 64-channel amplifier chips and one of the RHS2116 16-channel stimulator/amplifier chips, allowing for simultaneous recording from up to 528 channels and stimulation from up to 16 channels. In addition, each board allows for a hardware reference from one of 16 sites distributed across the array. The digitized data is transferred from the amplifier boards to an associated Intan Technologies 1,024-channel RHD controller or 126-channel RHS controller using low-voltage differential signaling (LVDS), where it is then stored on a USB-connected computer.

The amplifier boards are designed to allow each board to be easily coupled to any array-interposer assembly through the inclusion of an array of pogo pins that make contact with an associated pad on the array-interposer assembly, connecting each electrode site with an amplifier input. These two boards are aligned and held together by two plates with integrated alignment features placed on the outward-facing sides of the boards and screwed together. Additional protection of these electronics is provided by a custom, 3D-printed casing with strain-relief features for the electrode array and optional mounting braces to fix the entire assembly to the skull.

The 1024-channel configuration was similar to the above, but with 16 64-channel amplifier chips required for all of the recording electrodes, and with references and stimulation electrodes wired externally as needed. These recording boards were attached using mezzanine connectors rather than pogo pins, for a more miniaturized interface.

#### Recording Software and Data Pre-processing

The recording computers interface with either controller via a custom configuration of the Intan Technologies RHX Data Acquisition Software, which allows for real-time event-triggered averaging in addition to base functionality. The sampling rate for recording is set at 20 kHz per channel, generating data at a rate around 2 GB per minute for each set of 1,024 channels. A 60 Hz notch filter is applied online during recording. For post-hoc analysis of local field potentials, unless otherwise specified, data is first down-sampled to 5 kHz using a Fourier method, then processed with a 5th-order Butterworth low-pass filter at 250 Hz.

#### Software Methods

Data processing is performed in C/C++, Python, and MATLAB, using community standard frameworks, including but not limited to Qt, CUDA, NumPy, SciPy, PyTorch, and Matplotlib.

Machine learning model training is performed using PyTorch, accelerated by an NVIDIA RTX 4090. Machine learning model inference is performed using TorchScript, utilizing 12th generation Intel Core i7 processors.

Data is parsed in real-time by configuring Intan RHX to serialize data using the Intan DAT format. DAT format serialization reduces the latency between data acquisition on-chip, and serialization to disk, when compared to the default RHD format, which buffers and writes to disk in 128-sample chunks. This enables smooth and consistent playback, regardless of sampling frequency.

Real-time visualizations (Figure 3i) are implemented using the Qt framework. Visualization of amplifier data is either performed using raw data, with no post-processing applied, or with one or more filters applied. Most commonly used for real-time analysis was a simple kernel smoothing technique applied to minimize the visual effects of lower quality channels, or reduce the amount of low amplitude, per-channel noise, which can lead to flickering of channels. While a number of kernel smoothing techniques can be applied, the most commonly used is the 3×3 Gaussian kernel.

Trials are generated by reading and combining amplifier samples and digital input signals, sampled simultaneously. Amplifier data is buffered in memory and aligned with the corresponding digital input signals. When a TTL high signal is detected on a predefined channel of the digital input, a number of amplifier samples are selected preceding and following the digital signal, and emitted as a trial. Additional processing of trial data may be performed, depending on the experimental setup or analytical techniques being used downstream, although the methods vary depending on experimental setup and the downstream analytical techniques being applied.

Trials are combined with external metadata to classify each trial as associated with a particular stimulus site. For example, in the case of somatosensory evoked potentials in the Göttingen minipig, these stimuli may correspond to sites on the rostrum, or to the null case when no stimulus was applied. Model training follows the standard procedure for model training and evaluation in PyTorch or MATLAB. Sampled data is shuffled and partitioned into “testing” or “training” collections for each epoch of training. Following training, model inference is performed in line with subsequent trial generation by evaluating the model on each trial, as the trial is emitted.

#### Free Recording of Spontaneous Cortical Activity

Example spectrograms are generated from data obtained at 20 kHz per channel, where spectral density is computed for a temporal resolution of 50 ms and frequency resolution of 17.5 Hz using a Hann window.

To demonstrate spatial correlation between pairs of electrodes, 20 kHz stimulus-free neural data was separated into continuous 2 s segments. Within each segment, the squared Pearson correlation coefficient *r*^2^ is computed for each pair of electrodes and associated to the corresponding physical electrode distance. The average *r*^2^ values across an array are pooled across 60 segments and averaged for each electrode distance.

#### Evoked Potentials

Somatosensory evoked potentials (SSEPs) were evoked by applying periodic pressure on the rostrum or peripheral nerve electrical stimulation. SSEPs caused by rostrum stimulation (rostrum SSEP) are measured by manually applying pressure at six different locations on the rostrum using a conical tip. The onset of a stimulus is defined as the instance when 0.1 lbf of force is applied to the rostrum. For peripheral nerve SSEP, electrical stimulation was applied to median and tibial nerves of each side of the animal by placing twisted subdermal needle electrodes (13 mm–27 Gauge, Cadwell, Kennewick, Washington, United States of America) near the location of the nerves. Repetitive stimuli (300 uS pulse at 2.79 Hz, more than 300 times) were presented using a Cadwell Cascade IOMax System with a limb module at intensities 1.5–2 times the threshold required to visualize twitching in the muscle distal to the stimulated nerve. Each nerve was stimulated separately while cortical responses were recorded by the electrode array. Neural response waveforms were temporally aligned to the stimulus onset. SSEPs were then computed as the averaged time-aligned signals over 250 stimuli for peripheral nerve SSEP and 140 stimuli for rostrum SSEP.

To elicit visual evoked potentials (VEP), the eyelid corresponding to the stimulated retina was retracted temporarily while periodic 50 ms flashes were generated at 1 Hz from an array of white light-emitting diodes. Neural response waveforms were temporally aligned to the stimulus onset. VEPs were calculated as the time-aligned averaged signals over 150 trials.

#### Cortical Stimulation

Electrical stimulation at the cortical surface was applied at one of the 200 µm electrodes, controlled by the Intan Technologies RHS controller and RHX software. Charge-balanced, biphasic, cathodic-first, 200 µs pulses of 100 µA peak current were delivered at 0.25 Hz. The evoked potentials were recorded over a series of trials. During analysis, for each trial and electrode, the “activity” of each trial was computed as the variance of the signal from 200 ms to 2000 ms post-stimulation, and the average activity was taken over 40 trials.

#### Electrophysiologic Recording and Motion Capture During Awake Locomotor Activity

A 1024-channel array was placed over the sensorimotor cortex on each hemisphere following carefully sized bilateral craniectomies. Two Intan 1024-channel RHD controllers were used to record from both arrays simultaneously.

A harness (Ruffwear, Oregon, United States of America) was placed on the animal while it was on the operating table before being transported to the treadmill (Firepaw, Plovdiv, Bulgaria).

To capture motions of the animals, a pair of OAK-1 cameras (Luxonis, CO, USA) was used, with one camera placed on each side of the animal. One camera was to cover anterior view of the animal to capture head and forelimb movements and the other was used to capture a posterior view, including hindlimb movements. Videos were recorded at 60 frames per second using a modified set of codes from DepthAI SDK provided by the camera manufacturer. Each pair of cameras was synchronized frame-by-frame. In order to synchronize videos and neural recordings, 4 light-emitting diodes (LEDs) were placed on the treadmill and were controlled by an Arduino microcontroller that controlled the LEDs with a pulse whose “on” state duration was 100 ms at 0.25 Hz. At least one LED is captured in each of the videos on each side, and the pulse was recorded as a digital input at 20 kHz by the Intan controller.

After recording sessions, each video was annotated using Premier Pro 2023 (Adobe, CA, USA) to classify the animal behaviors into one of the following three states: resting, limb movements, and head movements. Limb movements included locomotion forward or backward, discrete one-limb movements, non-locomotive multiple-limb movements, or a sequence of those limb movements without any rest. A movement onset frame was defined to be a frame where one or more limbs started to move. During resting, none of the body parts visible in any of the videos moves. A beginning and an end of each class are annotated.

#### Human Intraoperative Recording

Human intraoperative recording, real-time analysis, and visualization were performed using 1024-electrode arrays configured with customized head stages and software configured as described in earlier sections, but packaged in a manner designed to facilitate ethylene oxide sterilization and secure fixation in the surgical field.

Spontaneous cortical activity and upper-limb somatosensory evoked potentials were obtained as described in earlier sections.

During awake language mapping, auditory cues provided by the examiner or visual cues (single words) presented on a screen instructed the patient to speak individual words. The cues and the full auditory output of the patient were recorded and time-synchronized with the electrophysiologic data for offline analysis.

### Neural Decoding

#### Sensory Decoding

Multi-class single-shot decoding efficacy is demonstrated by classifying array-wide neural recordings of rostrum SSEPs using a convolutional neural network (CNN). For each stimulation location, the stimulus was analog in space (within 3 mm radius from target), duration (200–400 ms), and applied force (0.2–0.6 lbf peak), reflecting the variance of real-world stimuli. The onset of a stimulus is defined as the moment when 0.1 lbf is applied to the rostrum. Recordings were first downsampled from 20 kHz to 1 kHz, and then low-pass filtered with a 490 Hz 5th-order Butterworth filter. Recording segments of 300 ms duration (125 ms pre-onset and 175 ms post-onset) were each associated with one of six stimulation locations or spontaneous activity, yielding seven classes. Each location was stimulated in 150–200 trials with an 80%-20% split into training and testing sets. The model was trained over 1500 iterations using cross entropy loss and gradient-descent ADAM optimizer. L1 regularization was employed for all weights.

The baseline convolutional neural network (Figure 4c) consists of two convolutional blocks followed by a 7-unit fully connected layer, leaky ReLU activation, another 7-unit fully connected layer, and finally a softmax layer. Each convolutional block carries out the following sequence: (1) filter with 3×3 kernels, stride 2, same padding, (2) leaky ReLU, (3) 2×2 max pooling with stride 2, (4) batch normalization, and (5) 50% dropout at training time. The first convolutional block has 25 filters, and the second block has 7. Decoding performance is presented as a confusion matrix showing predicted versus actual labels, normalized to the number of predicted labels per class.

Decoding with reduced electrode density or reduced spatial coverage is simulated by using only data collected from a subset of electrodes. For low electrode density, data from every other row and column of electrodes was used, which is equivalent to employing an electrode array with the same spatial coverage and one-fourth the density of the complete array. For low spatial coverage, a continuous rectangular region that is one-quarter the size of the whole array was chosen for the highest decoding accuracy for the greatest number of rostrum stimulation locations. This region is equivalent to an electrode array with one-quarter the spatial coverage and the same electrode density as the complete array. These two reduced electrode configurations yield the same amount of recording data as one another, but one-quarter that of the complete array. Consequently, the number of inputs and units in hidden layers for the convolutional neural network are adjusted accordingly, while enforcing seven output classes. A total of 50 models were trained for each electrode configuration: whole-array, one-fourth density, and one-fourth coverage. The average decoding accuracy and standard error of the mean for each stimulation location is presented.

A different CNN-based architecture (DenseSparseNet) was developed to improve decoding accuracy, allowing for the likelihood that correlations between nearby and distant electrodes may capture different types of useful information. This architecture splits the convolution steps into three streams, each with convolutional blocks applied to a grid of electrodes subsampled at different densities. Convolutional blocks are equivalent to those of the baseline CNN model but with max pooling removed, and each convolutional layer and stream has 5 filters. After each application of a convolutional block, the higher density streams are (1) passed forward to another convolutional block, and in parallel (2) subsampled spatially and then added to the stream with the next level of density (“recombination step”). The recombination step is carried out from highest to lowest density, unidirectionally in sequence. The two layers of convolution and recombination are followed by fully connected and activation layers analogous to those of the baseline CNN model.

#### Motor Decoding in Consciously Behaving Large Animals

For motor decoding, three classes of behavior were considered: head movements, limb movements, and rest. Movement events preceded by at least 750 ms of rest and the rest events that lasted at least 2 seconds were chosen. For one of the sessions, we included sensory stimulation of the rostrum and aligned the data to the onset of the stimulation. Micro-ECoG recording data was down-sampled from 20 kHz to 1 kHz and was aligned to the movement onset of limb and head movements or to the middle of resting periods to contain the data segment [-500 ms, 500 ms] around the alignment point. We then normalized the data for each μECoG channel across all trial classes to have zero mean and equal variance. To train on a balanced number of samples per behavior class, we employed undersampling by matching the number of samples in the minority behavior class. The first samples were selected to keep the effect of the anesthesia and the activity level of the animal consistent. For validation of decoding performance, *k*-fold cross validation was used (*k*=5 or 10) to compute accuracy statistics and confusion matrices.

Due to the smaller number of motor events in comparison to sensory stimulation events, we employed a simpler CNN architecture to decode the behavioral state of an animal in each trial to avoid overfitting. The input layer of the network receives per-μECoG-channel normalized neural signals from each trial as a data array having dimensions equal to the number of ECoG channels (2048) by the chosen number of time subintervals (40, corresponding to 40 25 ms averaging intervals per 1 s of data collected). The architecture comprises four consecutive 2D convolutional layers, each with increasing numbers of filters (8, 16, 32, 64) and a kernel size of 3×3 with “same” padding. After each convolutional layer, batch normalization and a ReLU activation function are applied, followed by a max-pooling layer with a pool size of 2×2 and a stride of 2. For the fourth convolutional layer, the max-pooling layer was omitted. The architecture concludes with a fully connected layer, a softmax layer, and a classification layer for each behavior state type. A cross-entropy loss function and a stochastic gradient descent algorithm with momentum optimization were used to train the network.

#### Speech Decoding from Patients Undergoing Awake Craniotomies for Language Mapping

We employed a logistic regression model on 4 minutes of neural data during which a patient spoke from a limited vocabulary of single-syllable words. The model was trained and tested on 61 trials with speech and 61 trials without speech. The 1024-channel neural data was selected in the time window of 1.5 seconds around the speech onset (0.5 second pre-speech and 1.0 second post-speech onset) and 1.5 seconds without speech during which the patient was resting. We then evaluated the model performance by running the prediction 100 times by randomly resampling the training and test data by 9:1 ratio without replacement.

## Author Contributions

Study conception and design: BIR, EH, MH, DP, AP, ML, MM, MM, CHM. Data collection: BIR, EH, MH, DP, ML, AP, KG, KT, SR, YWB, SHL, SB, PEK, CHM. Interpretation of results: BIR, EH, MH, DP, AP, MM, DT, MM, KR, TH, VT, YWB, CHM. Manuscript preparation: BIR, EH, MH, DP, MV, AP, MM, YWB, CHM. All authors reviewed the results and approved this version of the manuscript.

## Acknowledgments

The authors gratefully acknowledge technical assistance from Oleg Modik on issues related to clinical neurophysiology and intraoperative monitoring, and assistance with quality assurance and documentation from Ronald Gonterman. The authors also gratefully acknowledge clinical assistance from colleagues at the West Virginia University School of Medicine and Rockefeller Neuroscience Institute in support of intraoperative mapping studies in neurosurgical patients. The authors also gratefully acknowledge the assistance of Jason L. Chen, Jonathan Steinberger, and Rony Abovitz for critical review of this manuscript, and the artistic and logistical assistance of Helen Melville and Ariana Dimock.

Correspondence and requests for materials should be addressed to Benjamin I. Rapoport at.

## Supplementary Information

**Supplementary Figure 1:**
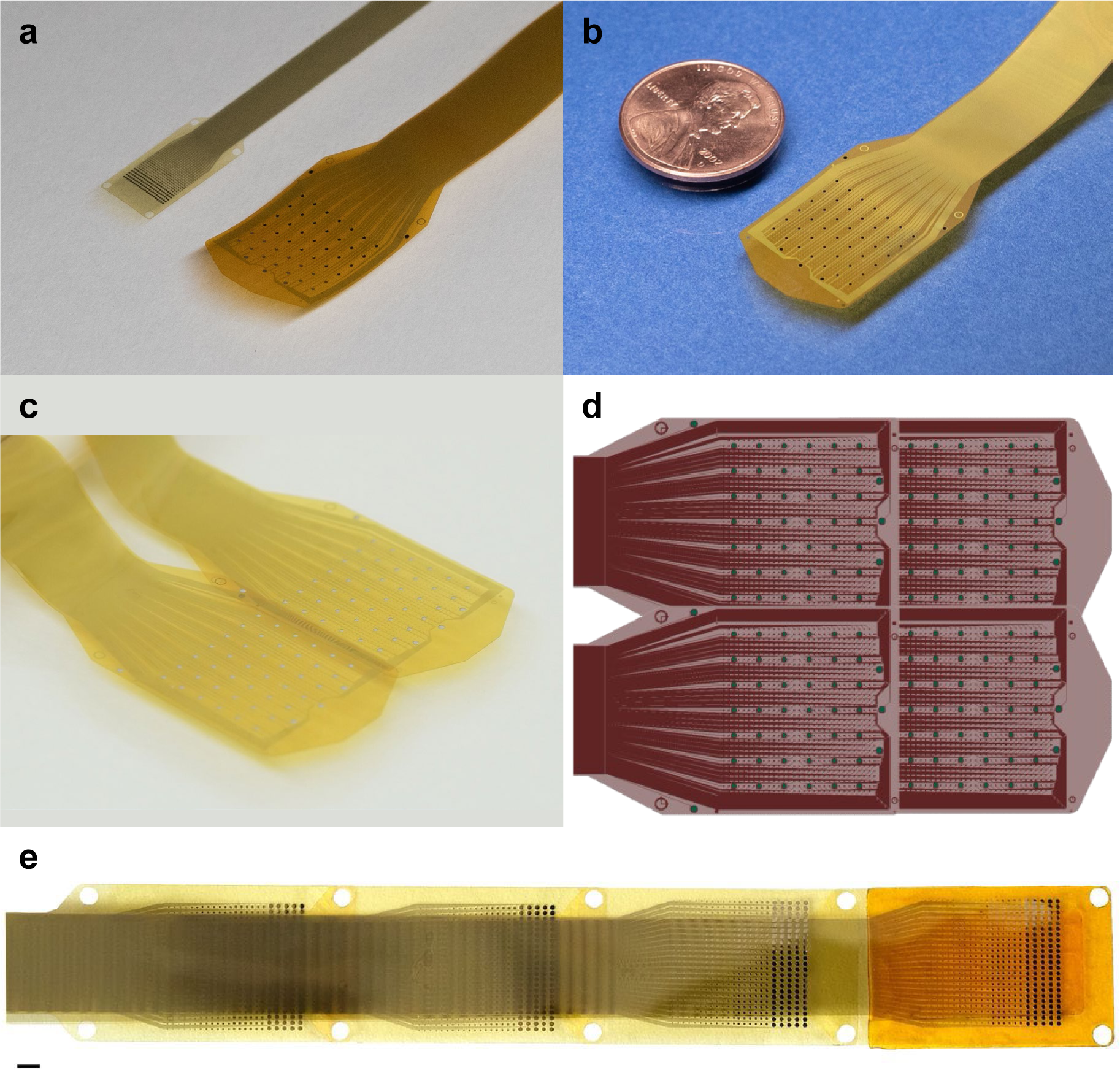
Individual and modular configurations of 1024-channel and 529-channel thin-film micro-electrocorticography arrays provide configurable options for cortical surface interfaces. (a) Device photographs of 529-channel (left) and 1024-channel (right) thin-film microelectrode arrays. (b) Photograph of 1024-channel thin-film microelectrode array with penny for scale. (c) Device photograph of two modularly connected 1024-channel thin-film microelectrode arrays. (d) Computer-aided design showing one possible modular (2-by-2) configuration of four 1024-channel arrays for simultaneous implantation of 4096 microelectrodes over 6.24 cm^2^ of cortical surface area. (e) Modular configurations of multiple 529-channel arrays can also be constructed. Shown here is a quadruply-connected assembly of four 529-electrode modules with a single pocket, for simultaneous implantation of 2116 microelectrodes over 1.92 cm^2^ of cortical surface area. Scale bar: 1 mm.

**Supplementary Figure 2.**
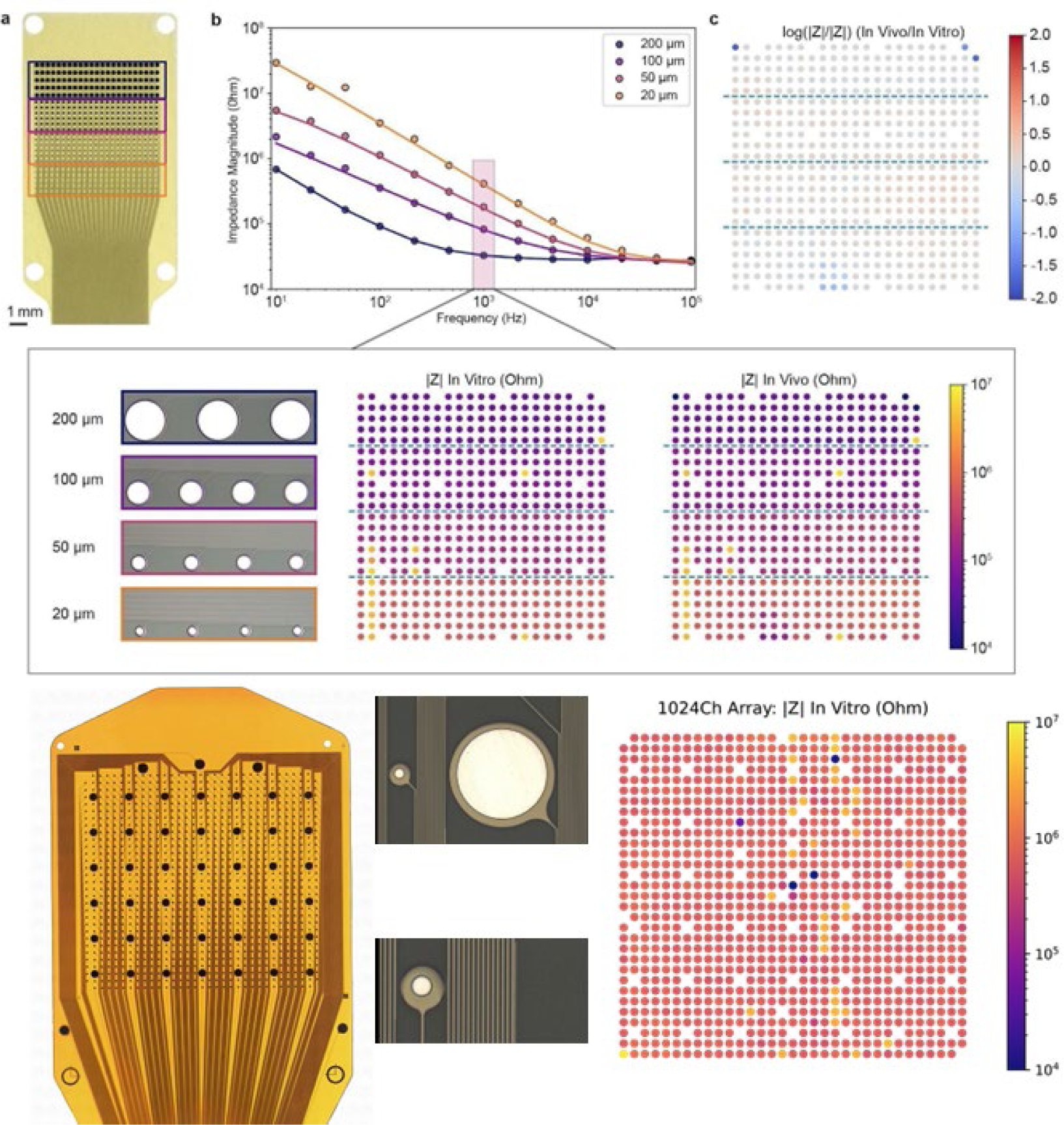
Electrode characterization. (a) Photograph of a single array with 200, 100, 50, and 20 µm electrodes outlined by blue, purple, pink, and orange boxes, respectively. (b) *In vitro* electrochemical impedance spectroscopy (EIS) spectra for each electrode size, with equivalent circuit fits based on a Randles circuit. (Inset) *In vitro* (left) and *in vivo* (right) impedance maps of recording channels over a complete 529-channel array, measured at 1 kHz, with microscope images for each electrode size included as references. Stimulation and ground channels are shown as gaps in the plots. (c) Map of the ratio between *in vivo* and *in vitro* impedance at 1 kHz, displaying minimal change across most of the array following implantation. (d) Photograph of a 1024-channel array configuration, with reference electrodes around the periphery and stimulation (large) and recording (small) electrodes, respectively, in the main rectangular cluster. (e) Micrographs showing detailed views of 380-micron stimulation (top) and 50-micron recording electrodes (bottom), with nearby trace routing also depicted. (f) Impedance map of recording electrodes only, measured at 1 kHz. (Stimulation electrodes are routed and measured separately from the record electrodes, as external pads, not pictured.)

**Supplementary Figure 3.**
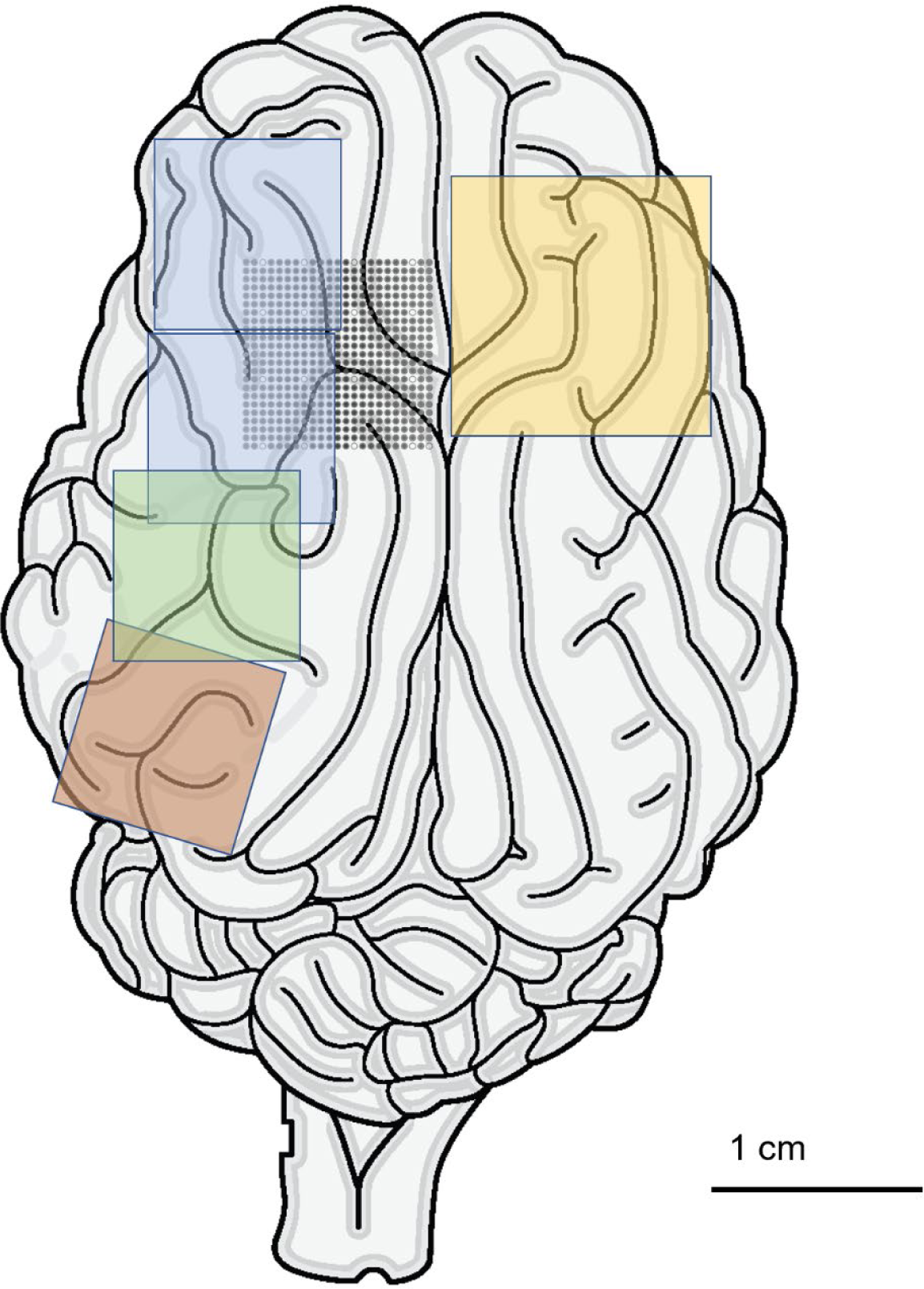
Cortical regions accessed in the Göttingen minipig using surface microelectrode arrays. Multiple functional regions of the Göttingen minipig brain were accessed in 26 animals during the course of these investigations. Array locations are shown here in schematic fashion. Although many experiments involved bilateral array placement, for clarity of illustration 529-channel array silhouettes are shown here over the left hemisphere, and a 1024-channel array is shown over the right hemisphere. Yellow, 1024-channel array over rostrum somatosensory and limb, face, and mouth motor regions. Blue, 529-channel arrays over rostrum somatosensory regions; green, 529-channel array over auditory cortex; brown, 529-channel array over visual cortex. Black dots, 529-channel array over limb, face, and mouth motor regions, each dot representing a single recording electrode.

**Supplementary Figure 4.**
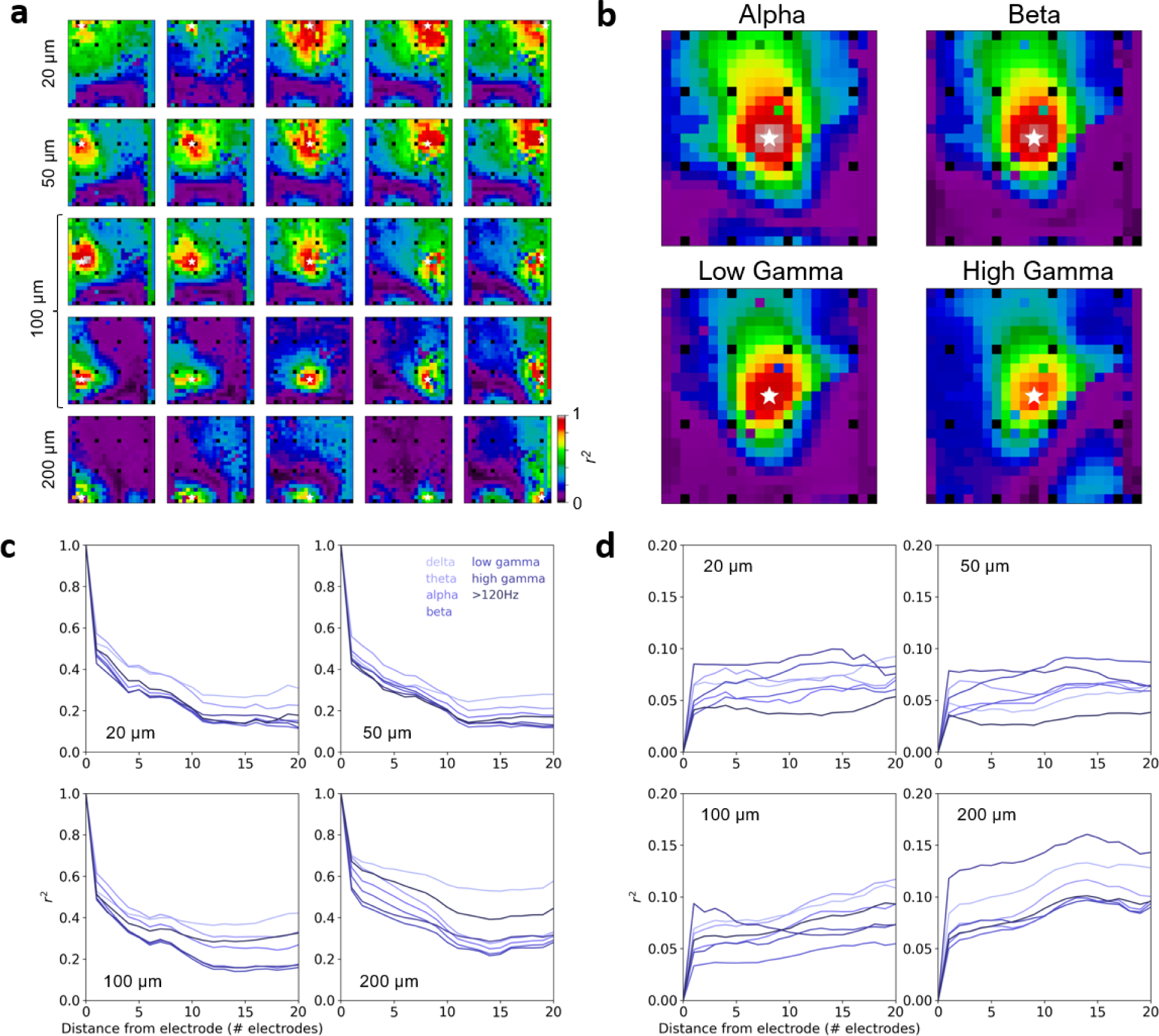
Spatial scales of information representation on the cortical surface. The degree to which closely spaced microelectrodes across a single 529-channel array record correlated or uncorrelated information is relevant to neural decoding and is illustrated here in several ways. (a) Correlation map of the Pearson correlation coefficient *r*^2^ computed for signals recorded during spontaneous cortical activity, with correlations across the entire array in each sub-plot referenced to one of 25 electrodes (white stars) distributed evenly over the array, as represented by the array of subplots. (b) Correlation map for different EEG bands recorded on a representative electrode. Low-gamma band is defined as 30-55Hz, and high gamma band is defined as 65-120Hz. (c) The dependence of correlation on electrode separation as a function of electrode size and EEG band. (d) Standard deviation corresponding to (c).

**Supplementary Figure 5.**
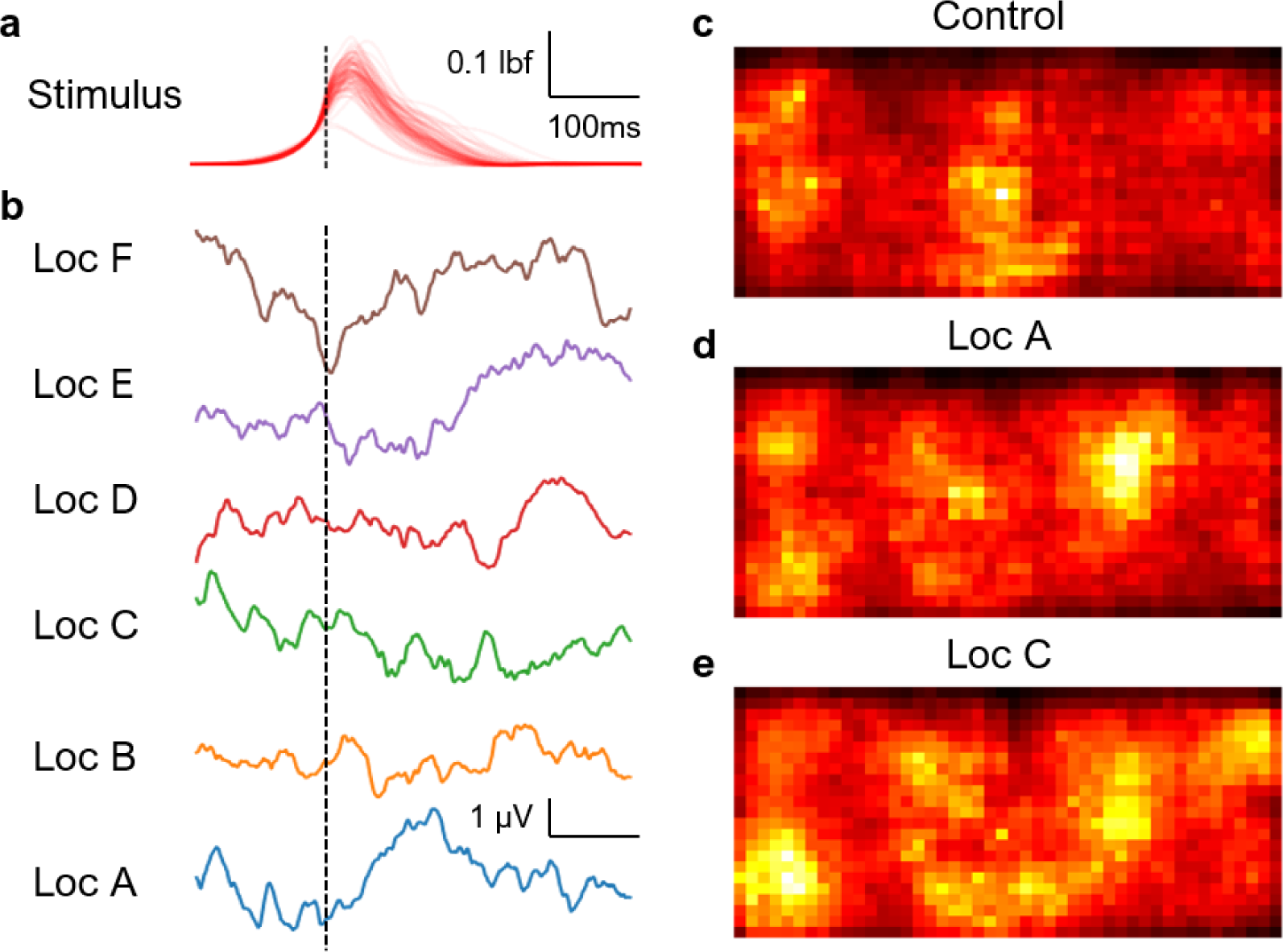
Saliency maps. (a) Force profile of 100 tactile stimuli (1 red line per trial) applied to the rostrum. All trials were time aligned to reaching 0.1 lbf of the rising edge. (b) Rostrum somatosensory evoked potential (SSEP) averaged over 130 trials for each stimulation location as shown in Figure 4a. The right column shows examples of average saliency maps for decoding single-shot (c) spontaneous trials (control), (d) stimulation at location A and (e) location C. The color scale is normalized to the maximum value for each stimulation location independently.

**Supplementary Figure 6.**
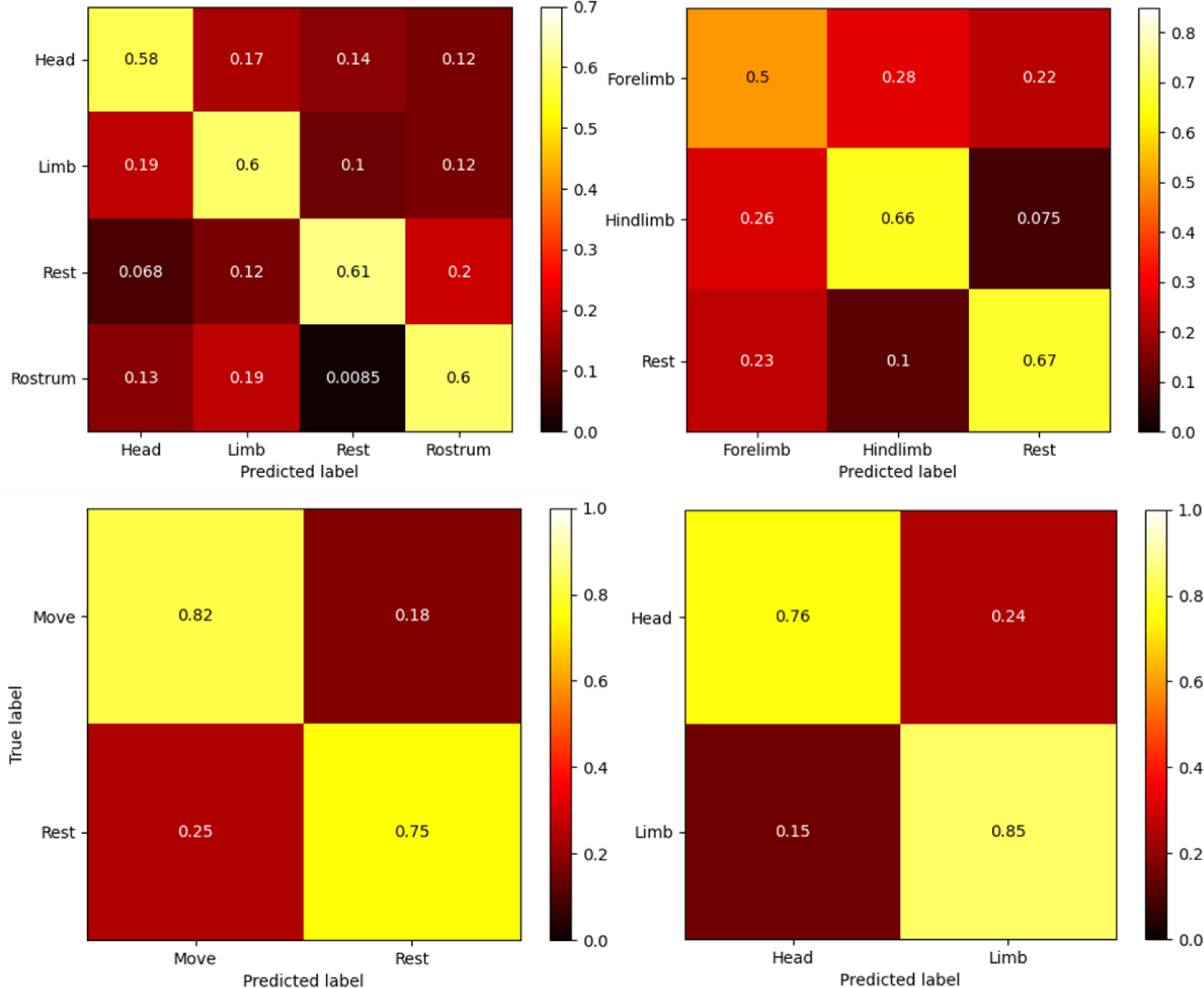
Versatile motor decoding paradigms from sensorimotor region. Confusion matrices from decoding sensory and motor modalities of various body regions (*k*-fold cross-validation; *k*=5). (a) Classification of head movement, limb movement, rest, and sensory stimulation of the rostrum (Accuracy: 60%). (b) Classification of forelimb movement, hindlimb movement, and rest (Accuracy: 61%). (c, d) Classification of general movement versus rest (Accuracy: 79%) and head versus limb movements (Accuracy: 80%).

**Supplementary Figure 7.**
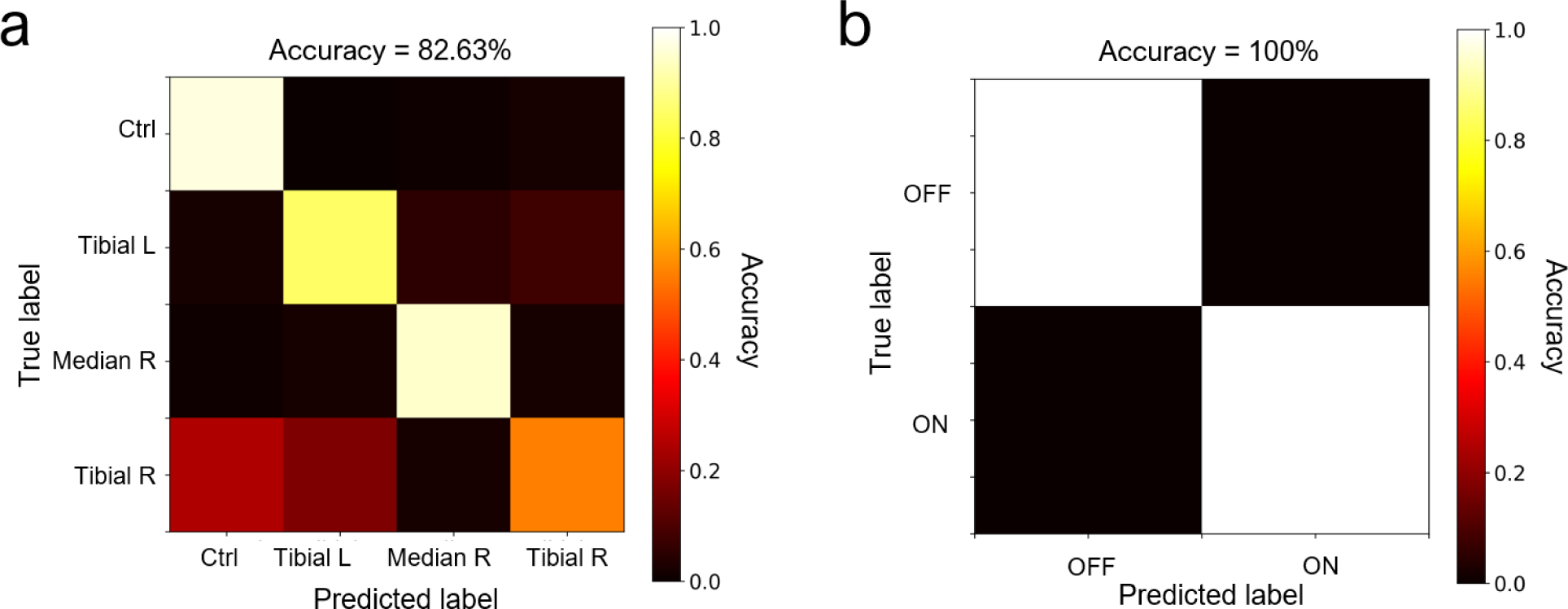
Multimodal neural decoding of somatosensory and visual information. (a) Confusion matrix for decoding somatosensory evoked potentials (SSEPs) induced through electrical stimulation of the median and tibial nerves (*R*, right; *L*, left. (b) Confusion matrix for decoding light flashes from the visual cortex as presented to the contralateral eye.

**Supplementary Video 1:**
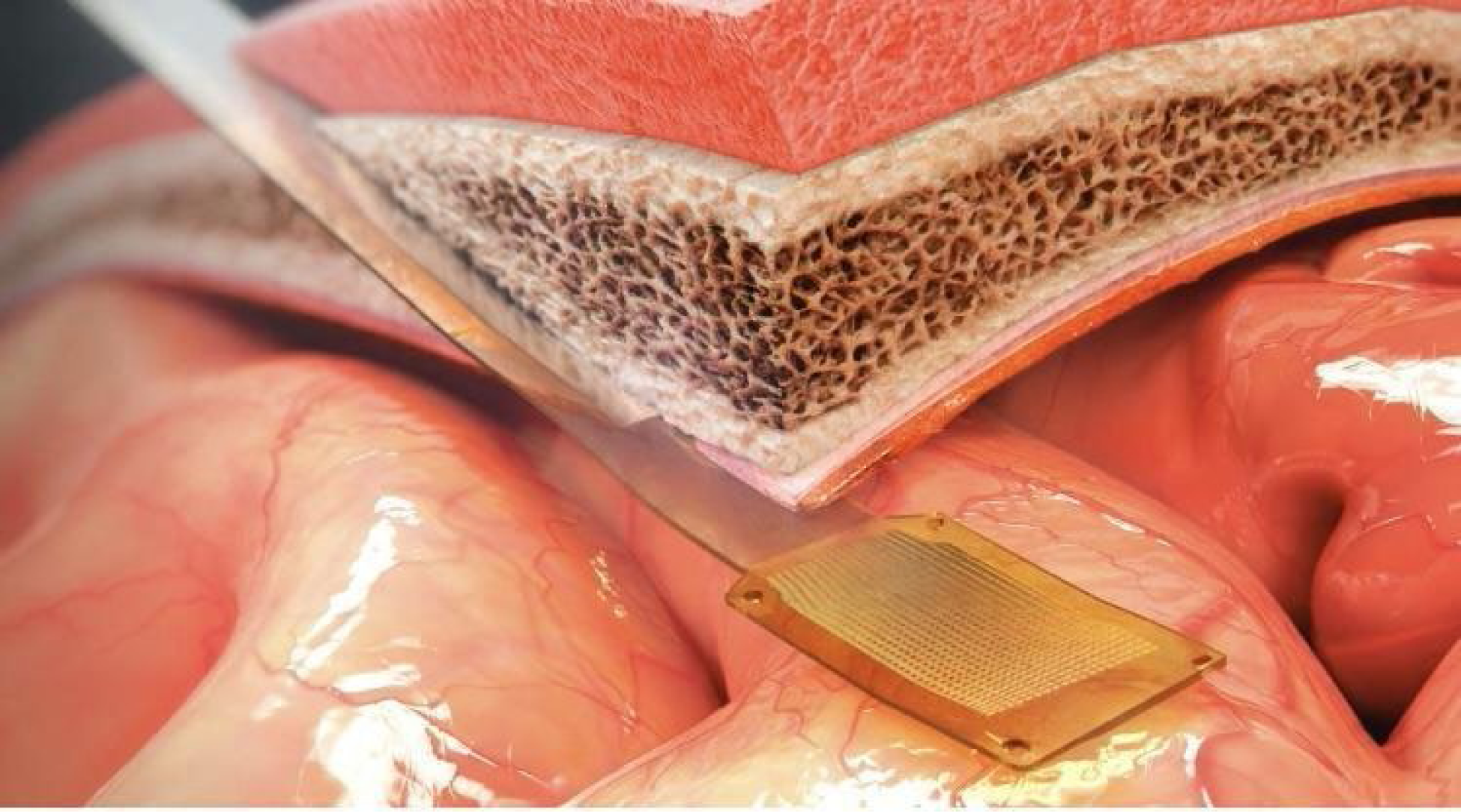
Cranial microslit insertion technique for placement of thin-film microelectrode arrays. This video demonstrates the cranial microslit insertion technique. The first segment illustrates forming the microslit at an approach angle tangent to the cortical surface target using a sagittal saw, first in the setting of an intact skull, then in a cut-away view. The next segment demonstrates the technique being employed *in vivo* in a Göttingen minipig. The final segment demonstrates the cranial microslit technique being used to perform subdural insertion of a 1024-channel microelectrode array in a human cadaver. (<u>Link to Supplementary Video 1</u>.)

**Supplementary Video 2:**
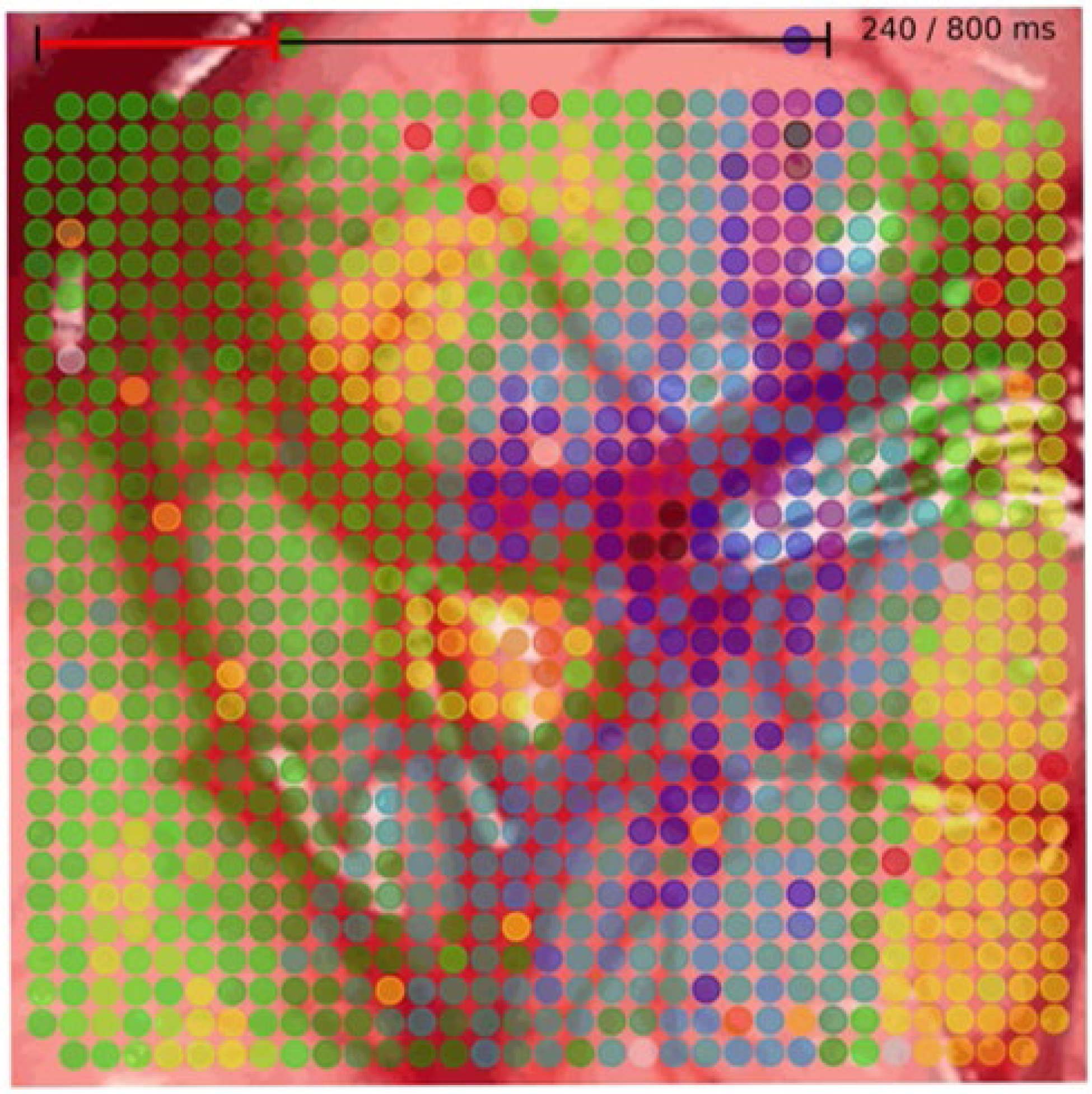
Thin-film microelectrode arrays provide high-resolution neural recordings from the cortical surface in real time under anesthesia. After implantation of a 1024-channel microelectrode array in a Göttingen minipig, neural recordings were obtained in the absence of any external stimulation, representing spontaneous electrocortical activity, as described in the manuscript. The video shows full-bandwidth neural recordings from one 1024-channel electrode array (right hemisphere) as a color-map of voltage, superimposed on and aligned to an image of the underlying cortical surface (each dot corresponds to one microelectrode in the array). (<u>Link to Supplementary Video 2</u>.)

**Supplementary Video 3:**
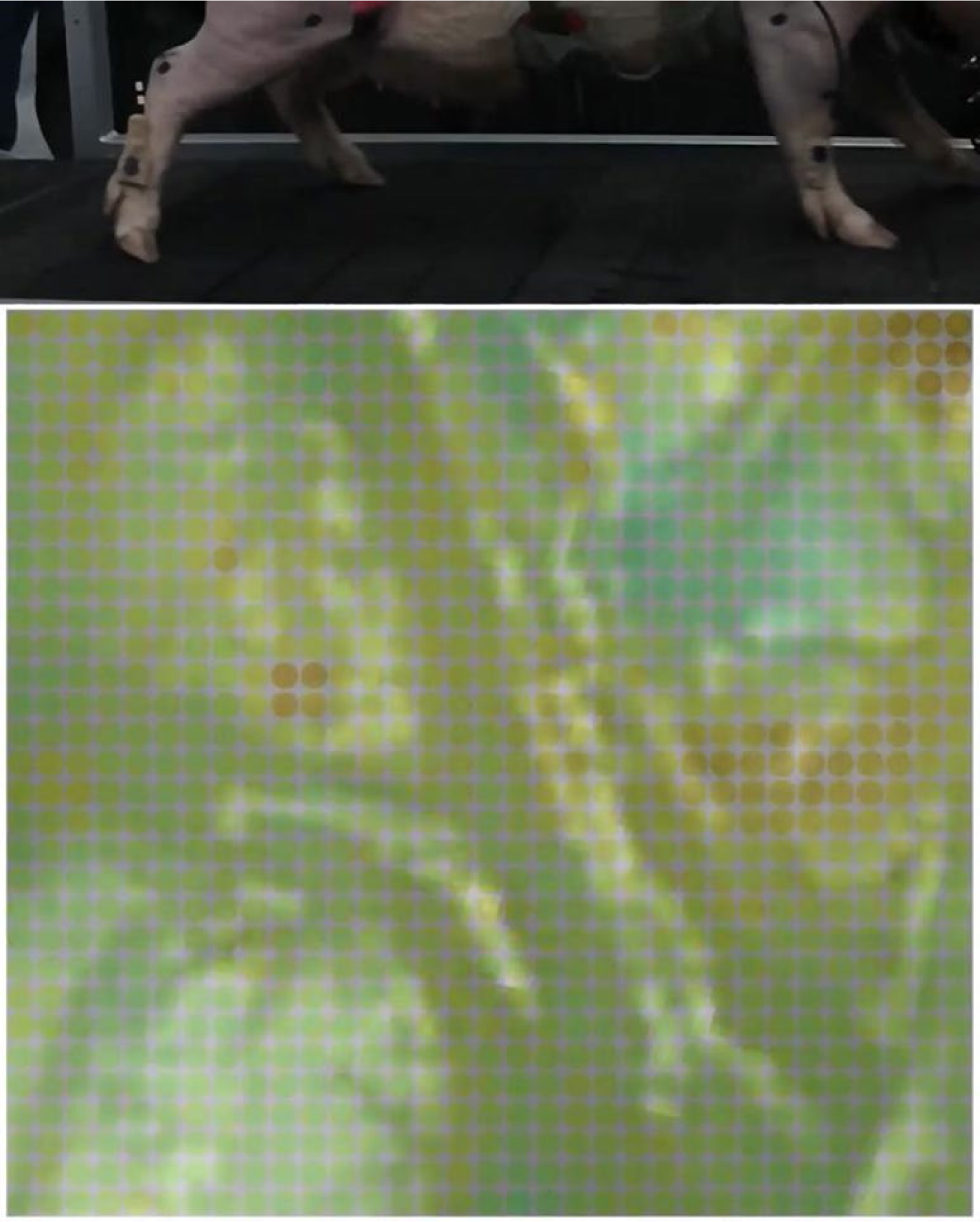
Thin-film microelectrode arrays provide high-resolution neural recordings from the cortical surface in real time during volitional walking. After bilateral implantation of two 1024-channel microelectrode arrays (one per hemisphere over sensorimotor cortex in a Göttingen minipig), the animal was awakened and permitted to walk ad lib on a treadmill. Head- and limb-based accelerometers and real-time motion tracking image capture data were time-synchronized with full-bandwidth neural recordings, and these data were used for neural decoding of volitional limb movement, as described in the manuscript. The video shows full-bandwidth neural recordings from one 1024-channel electrode array (left hemisphere) as a color-map of voltage, superimposed on and aligned to an image of the underlying cortical surface (each dot corresponds to one microelectrode in the array); limb movement during treadmill walking is shown in the top panel, time-aligned to the displayed neural recording. (<u>Link to Supplementary Video 3</u>.)

**Supplementary Video 4:**
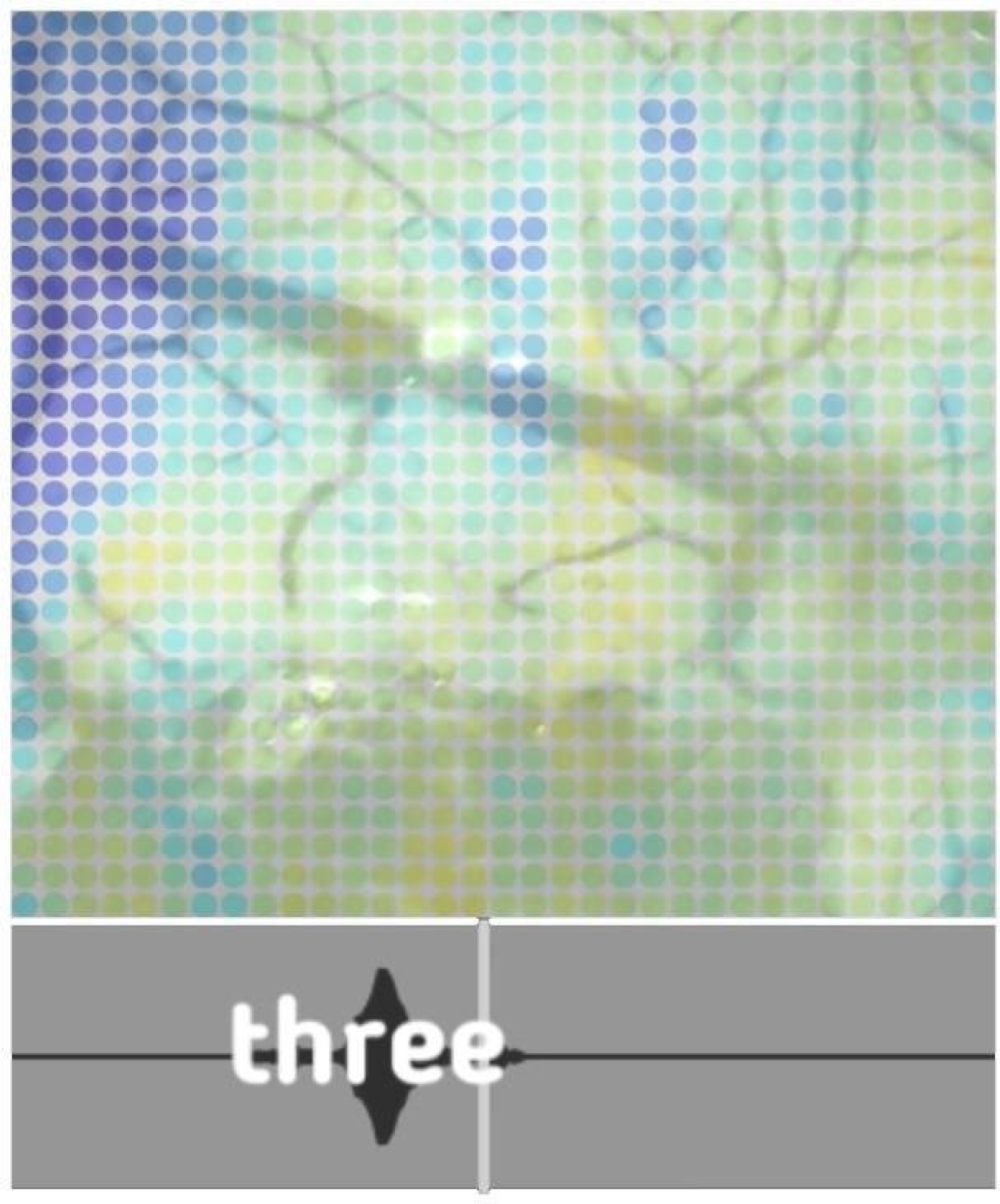
High-resolution neural recordings from the cortical surface during speech. Thin-film microelectrode arrays provide high-resolution neural recordings in real time during awake language mapping in a human neurosurgical patient. The video demonstrates the final phase of conventional cortical mapping (monopolar stimulation) followed by placement of the 1024-channel thim-film microelectrode array. Following array placement, the video shows a digital overlay-representation of electrocortical activity superimposed on an image of the cortical surface. Each colored disc represents a microelectrode placed in the corresponding location. The time-evolving color map represents normalized raw voltage as obtained from the digital steps of the analog-to-digital converter (*dark blue*, low; *yellow*, high). The time-synchronized audio amplitude of the cued words being spoken is shown in the lower panel, with superimposed textual annotation of the spoken words. (<u>Link to Supplementary Video 4</u>.)

## References

1. Bouton, C. E. et al. Restoring cortical control of functional movement in a human with quadriplegia. Nature 533, 247–250 (2016).

2. Kim, S.-P., Simeral, J. D., Hochberg, L. R., Donoghue, J. P. & Black, M. J. Neural control of computer cursor velocity by decoding motor cortical spiking activity in humans with tetraplegia. J. Neural Eng. 5, 455–476 (2008).

3. Pandarinath, C. et al. High performance communication by people with paralysis using an intracortical brain-computer interface. eLife 6, e18554 (2017).

4. Simeral, J. D., Kim, S.-P., Black, M. J., Donoghue, J. P. & Hochberg, L. R. Neural control of cursor trajectory and click by a human with tetraplegia 1000 days after implant of an intracortical microelectrode array. J. Neural Eng. 8, 025027 (2011).

5. Gilja, V. et al. Clinical translation of a high-performance neural prosthesis. Nat Med 21, 1142–1145 (2015).

6. Moses, D. A. et al. Neuroprosthesis for Decoding Speech in a Paralyzed Person with Anarthria. N Engl J Med 385, 217–227 (2021).

7. Wang, W. et al. An Electrocorticographic Brain Interface in an Individual with Tetraplegia. PLoS ONE 8, e55344 (2013).

8. Hochberg, L. R. et al. Neuronal ensemble control of prosthetic devices by a human with tetraplegia. Nature 442, 164–171 (2006).

9. Velliste, M., Perel, S., Spalding, M. C., Whitford, A. S. & Schwartz, A. B. Cortical control of a prosthetic arm for self-feeding. Nature 453, 1098–1101 (2008).

10. Schwartz, A. B., Cui, X. T., Weber, D. J. & Moran, D. W. Brain-Controlled Interfaces: Movement Restoration with Neural Prosthetics. Neuron 52, 205–220 (2006).

11. Collinger, J. L. et al. High-performance neuroprosthetic control by an individual with tetraplegia. The Lancet 381, 557–564 (2013).

12. Brindley, G. S. & Lewin, W. S. The sensations produced by electrical stimulation of the visual cortex. The Journal of Physiology 196, 479–493 (1968).

13. Dobelle, W. H., Mladejovsky, M. G. & Girvin, J. P. Artificial Vision for the Blind: Electrical Stimulation of Visual Cortex Offers Hope for a Functional Prosthesis. Science 183, 440–444 (1974).

14. Dobelle, W. H. et al. Artificial Vision for the Blind by Electrical Stimulation of the Visual Cortex. Neurosurgery 5, 521–527 (1979).

15. Eddington, D. K. Speech discrimination in deaf subjects with cochlear implants. The Journal of the Acoustical Society of America 68, 885–891 (1980).

16. Wilson, B. S. & Dorman, M. F. Cochlear implants: A remarkable past and a brilliant future. Hearing Research 242, 3–21 (2008).

17. Weiland, J. D., Liu, W. & Humayun, M. S. Retinal Prosthesis. Annu. Rev. Biomed. Eng. 7, 361–401 (2005).

18. Lewis, P. M., Ackland, H. M., Lowery, A. J. & Rosenfeld, J. V. Restoration of vision in blind individuals using bionic devices: A review with a focus on cortical visual prostheses. Brain Research 1595, 51–73 (2015).

19. Metzger, S. L. et al. A high-performance neuroprosthesis for speech decoding and avatar control. Nature 620, 1037–1046 (2023).

20. Willett, F. R. et al. A high-performance speech neuroprosthesis. Nature 620, 1031–1036 (2023).

21. Lorach, H. et al. Walking naturally after spinal cord injury using a brain–spine interface. Nature 618, 126–133 (2023).

22. Wang, W. et al. Human Motor Cortical Activity Recorded with Micro-ECoG Electrodes During Individual Finger Movements. Conf Proc IEEE Eng Med Biol Soc 2009, 586–589 (2009).

23. Wessberg, J. et al. Real-time prediction of hand trajectory by ensembles of cortical neurons in primates. Nature 408, 361–365 (2000).

24. Carmena, J. M. et al. Learning to Control a Brain–Machine Interface for Reaching and Grasping by Primates. PLoS Biol 1, e42 (2003).

25. Lebedev, M. A. & Nicolelis, M. A. L. Brain–machine interfaces: past, present and future. Trends in Neurosciences 29, 536–546 (2006).

26. Fraser, G. W., Chase, S. M., Whitford, A. & Schwartz, A. B. Control of a brain–computer interface without spike sorting. J. Neural Eng. 6, 055004 (2009).

27. Gilja, V. et al. A high-performance neural prosthesis enabled by control algorithm design. Nat Neurosci 15, 1752–1757 (2012).

28. Pandarinath, C. et al. Inferring single-trial neural population dynamics using sequential auto-encoders. Nat Methods 15, 805–815 (2018).

29. Collinger, J. L., Gaunt, R. A. & Schwartz, A. B. Progress towards restoring upper limb movement and sensation through intracortical brain-computer interfaces. Current Opinion in Biomedical Engineering 8, 84–92 (2018).

30. Musk, E. & Neuralink. An Integrated Brain-Machine Interface Platform With Thousands of Channels. J Med Internet Res 21, e16194 (2019).

31. Sahasrabuddhe, K., et al. The Argo: A 65,536 channel recording system for high density neural recording in vivo. http://biorxiv.org/lookup/doi/10.1101/2020.07.17.209403<x> (2020) doi:10.1101/2020.07.17.209403.

32. Schalk, G. et al. Decoding two-dimensional movement trajectories using electrocorticographic signals in humans. J. Neural Eng. 4, 264–275 (2007).

33. Wissel, T. et al. Hidden Markov model and support vector machine based decoding of finger movements using electrocorticography. J. Neural Eng. 10, 056020 (2013).

34. Moses, D. A., Leonard, M. K., Makin, J. G. & Chang, E. F. Real-time decoding of question-and-answer speech dialogue using human cortical activity. Nature Communications 10, 3096 (2019).

35. Sun, P., Anumanchipalli, G. K. & Chang, E. F. Brain2Char: A Deep Architecture for Decoding Text from Brain Recordings. Preprint at http://arxiv.org/abs/1909.01401 (2019).

36. Yu, B. M. et al. Mixture of Trajectory Models for Neural Decoding of Goal-Directed Movements. Journal of Neurophysiology 97, 3763–3780 (2007).

37. Kemere, C., Shenoy, K. V. & Meng, T. H. Model-Based Neural Decoding of Reaching Movements: A Maximum Likelihood Approach. IEEE Trans. Biomed. Eng. 51, 925–932 (2004).

38. Wilson, G. H. et al. Decoding spoken English from intracortical electrode arrays in dorsal precentral gyrus. J. Neural Eng. 17, 066007 (2020).

39. Harrison, R. R. & Charles, C. A low-power low-noise cmos for amplifier neural recording applications. IEEE J. Solid-State Circuits 38, 958–965 (2003).

40. Harrison, R. R. et al. A Low-Power Integrated Circuit for a Wireless 100-Electrode Neural Recording System. IEEE J. Solid-State Circuits 42, 123–133 (2007).

41. Sarpeshkar, R. et al. Low-Power Circuits for Brain–Machine Interfaces. IEEE Trans. Biomed. Circuits Syst. 2, 173–183 (2008).

42. Mora Lopez, C., et al. A Neural Probe With Up to 966 Electrodes and Up to 384 Configurable Channels in 0.13 $\mu$m SOI CMOS. IEEE Trans. Biomed. Circuits Syst. 11, 510–522 (2017).

43. Wang, S. et al. A Compact Quad-Shank CMOS Neural Probe With 5,120 Addressable Recording Sites and 384 Fully Differential Parallel Channels. IEEE Trans. Biomed. Circuits Syst. 13, 1625–1634 (2019).

44. Biederman, W. et al. A 4.78 mm 2 Fully-Integrated Neuromodulation SoC Combining 64 Acquisition Channels With Digital Compression and Simultaneous Dual Stimulation. IEEE J. Solid-State Circuits 50, 1038–1047 (2015).

45. Yoon, D.-Y. et al. A 1024-Channel Simultaneous Recording Neural SoC with Stimulation and Real-Time Spike Detection. in 2021 Symposium on VLSI Circuits 1–2 (IEEE, 2021). doi:10.23919/VLSICircuits52068.2021.9492480.

46. Johnson, B. C. et al. An implantable 700μW 64-channel neuromodulation IC for simultaneous recording and stimulation with rapid artifact recovery. in 2017 Symposium on VLSI Circuits C48–C49 (IEEE, 2017). doi:10.23919/VLSIC.2017.8008543.

47. Lee, J. et al. Neural recording and stimulation using wireless networks of microimplants. Nat Electron 4, 604–614 (2021).

48. Ha, S. et al. Silicon-Integrated High-Density Electrocortical Interfaces. Proc. IEEE 105, 11–33 (2017).

49. Ahmadi, N. et al. Towards a Distributed, Chronically-Implantable Neural Interface. in 2019 9th International IEEE/EMBS Conference on Neural Engineering (NER) 719–724 (IEEE, 2019). doi:10.1109/NER.2019.8716998.

50. Seo, D. et al. Wireless Recording in the Peripheral Nervous System with Ultrasonic Neural Dust. Neuron 91, 529–539 (2016).

51. O’Leary, G., Groppe, D. M., Valiante, T. A., Verma, N. & Genov, R. NURIP: Neural Interface Processor for Brain-State Classification and Programmable-Waveform Neurostimulation. IEEE J. Solid-State Circuits 53, 3150–3162 (2018).

52. Cogan, S. F. Neural Stimulation and Recording Electrodes. Annu. Rev. Biomed. Eng. 10, 275–309 (2008).

53. Wellman, S. M. et al. A Materials Roadmap to Functional Neural Interface Design. Adv. Funct. Mater. 28, 1701269 (2018).

54. Ordonez, J. S., Boehler, C., Schuettler, M. & Stieglitz, T. Improved polyimide thin-film electrodes for neural implants. in 2012 Annual International Conference of the IEEE Engineering in Medicine and Biology Society 5134–5137 (IEEE, 2012). doi:10.1109/EMBC.2012.6347149.

55. Vomero, M. et al. Incorporation of Silicon Carbide and Diamond-Like Carbon as Adhesion Promoters Improves In Vitro and In Vivo Stability of Thin-Film Glassy Carbon Electrocorticography Arrays. Adv. Biosys. 2, 1700081 (2018).

56. Deku, F. et al. Amorphous silicon carbide ultramicroelectrode arrays for neural stimulation and recording. J. Neural Eng. 15, 016007 (2018).

57. Li et al. Ultra-Long-Term Reliable Encapsulation Using an Atomic Layer Deposited HfO2/Al2O3/HfO2 Triple-Interlayer for Biomedical Implants. Coatings 9, 579 (2019).

58. Jeong, J. et al. Conformal Hermetic Sealing of Wireless Microelectronic Implantable Chiplets by Multilayered Atomic Layer Deposition (ALD). Adv. Funct. Mater. 29, 1806440 (2019).

59. Lamont, C. et al. Silicone encapsulation of thin-film SiO _x_, SiO _x_ N _y_ and SiC for modern electronic medical implants: a comparative long-term ageing study. J. Neural Eng. 18, 055003 (2021).

60. Cook, M. J. et al. Prediction of seizure likelihood with a long-term, implanted seizure advisory system in patients with drug-resistant epilepsy: a first-in-man study. The Lancet Neurology 12, 563–571 (2013).

61. Smalley, E. The business of brain–computer interfaces. Nat Biotechnol 37, 978–982 (2019).

62. Lebedev, M. A. & Nicolelis, M. A. L. Brain-Machine Interfaces: From Basic Science to Neuroprostheses and Neurorehabilitation. Physiological Reviews 97, 767–837 (2017).

63. Kang, Y. et al. Epidemiology of worldwide spinal cord injury: a literature review. JN 6, 1–9 (2017).

64. Lo, J., Chan, L. & Flynn, S. A Systematic Review of the Incidence, Prevalence, Costs, and Activity and Work Limitations of Amputation, Osteoarthritis, Rheumatoid Arthritis, Back Pain, Multiple Sclerosis, Spinal Cord Injury, Stroke, and Traumatic Brain Injury in the United States: A 2019 Update. Archives of Physical Medicine and Rehabilitation 102, 115–131 (2021).

65. Katan, M. & Luft, A. Global Burden of Stroke. Semin Neurol 38, 208–211 (2018).

66. Zack, M. M. & Kobau, R. National and State Estimates of the Numbers of Adults and Children with Active Epilepsy — United States, 2015. MMWR Morb. Mortal. Wkly. Rep. 66, 821–825 (2017).

67. Chan, T., Friedman, D. S., Bradley, C. & Massof, R. Estimates of Incidence and Prevalence of Visual Impairment, Low Vision, and Blindness in the United States. JAMA Ophthalmol 136, 12 (2018).

68. Matthews, K. A. et al. Racial and ethnic estimates of Alzheimer’s disease and related dementias in the United States (2015–2060) in adults aged ≥65 years. Alzheimer’s & Dementia 15, 17–24 (2019).

69. Vomero, M. et al. Conformable polyimide-based μECoGs: Bringing the electrodes closer to the signal source. Biomaterials 255, 120178 (2020).

70. Bockhorst, T. et al. Synchrony surfacing: Epicortical recording of correlated action potentials. Eur J Neurosci 48, 3583–3596 (2018).

71. Minev, I. R. et al. Electronic dura mater for long-term multimodal neural interfaces. Science 347, 159–163 (2015).

72. Viventi, J. et al. Flexible, foldable, actively multiplexed, high-density electrode array for mapping brain activity in vivo. Nat Neurosci 14, 1599–1605 (2011).

73. Khodagholy, D. et al. NeuroGrid: recording action potentials from the surface of the brain. Nat Neurosci 18, 310–315 (2015).

74. Khodagholy, D., et al. Highly Conformable Conducting Polymer Electrodes for In Vivo Recordings. Adv. Mater. 23, H268–H272 (2011).

75. Chung, J. E. et al. High-Density, Long-Lasting, and Multi-region Electrophysiological Recordings Using Polymer Electrode Arrays. Neuron 101, 21–31.e5 (2019).

76. Hirschberg, A. W. et al. Development of an anatomically conformal parylene neural probe array for multi-region hippocampal recordings. in 2017 IEEE 30th International Conference on Micro Electro Mechanical Systems (MEMS) 129–132 (IEEE, 2017). doi:10.1109/MEMSYS.2017.7863357.

77. Hara, S. A. et al. Long-term stability of intracortical recordings using perforated and arrayed Parylene sheath electrodes. J. Neural Eng. 13, 066020 (2016).

78. Kim, B. J. et al. 3D Parylene sheath neural probe for chronic recordings. J. Neural Eng. 10, 045002 (2013).

79. Chiang, C.-H. et al. Flexible, high-resolution thin-film electrodes for human and animal neural research. J. Neural Eng. 18, 045009 (2021).

80. Duraivel, S. et al. High-resolution neural recordings improve the accuracy of speech decoding. Nat Commun 14, 6938 (2023).

81. Kim, D.-H., Ghaffari, R., Lu, N. & Rogers, J. A. Flexible and Stretchable Electronics for Biointegrated Devices. Annu. Rev. Biomed. Eng. 14, 113–128 (2012).

82. Rubehn, B., Bosman, C., Oostenveld, R., Fries, P. & Stieglitz, T. A MEMS-based flexible multichannel ECoG-electrode array. J. Neural Eng. 6, 036003 (2009).

83. Salari, E. et al. Classification of Articulator Movements and Movement Direction from Sensorimotor Cortex Activity. Sci Rep 9, 14165 (2019).

84. Schalk, G. et al. Two-dimensional movement control using electrocorticographic signals in humans. J. Neural Eng. 5, 75–84 (2008).

85. Pistohl, T., Ball, T., Schulze-Bonhage, A., Aertsen, A. & Mehring, C. Prediction of arm movement trajectories from ECoG-recordings in humans. Journal of Neuroscience Methods 167, 105–114 (2008).

86. Anumanchipalli, G. K., Chartier, J. & Chang, E. F. Speech synthesis from neural decoding of spoken sentences. Nature 568, 493–498 (2019).

87. Angrick, M. et al. Speech synthesis from ECoG using densely connected 3D convolutional neural networks. J. Neural Eng. 16, 036019 (2019).

88. Ho, E., et al. The Layer 7 Cortical Interface: A Scalable and Minimally Invasive Brain– Computer Interface Platform. bioRxiv 2022.01.02.474656 (2022) doi:10.1101/2022.01.02.474656.

89. Polikov, V. S., Tresco, P. A. & Reichert, W. M. Response of brain tissue to chronically implanted neural electrodes. Journal of Neuroscience Methods 148, 1–18 (2005).

90. Biran, R., Martin, D. C. & Tresco, P. A. Neuronal cell loss accompanies the brain tissue response to chronically implanted silicon microelectrode arrays. Experimental Neurology 195, 115–126 (2005).

91. Winslow, B. D. & Tresco, P. A. Quantitative analysis of the tissue response to chronically implanted microwire electrodes in rat cortex. Biomaterials 31, 1558–1567 (2010).

92. Volkova, K., Lebedev, M. A., Kaplan, A. & Ossadtchi, A. Decoding Movement From Electrocorticographic Activity: A Review. Front. Neuroinform. 13, 74 (2019).

93. Branco, M. P. et al. Decoding hand gestures from primary somatosensory cortex using high-density ECoG. NeuroImage 147, 130–142 (2017).

94. Nakanishi, Y. et al. Prediction of Three-Dimensional Arm Trajectories Based on ECoG Signals Recorded from Human Sensorimotor Cortex. PLoS ONE 8, e72085 (2013).

95. Pistohl, T. et al. Grasp Detection from Human ECoG during Natural Reach-to-Grasp Movements. PLoS ONE 8, e54658 (2013).

96. Pistohl, T., Schulze-Bonhage, A., Aertsen, A., Mehring, C. & Ball, T. Decoding natural grasp types from human ECoG. NeuroImage 59, 248–260 (2012).

97. Willett, F. R. et al. A high-performance speech neuroprosthesis. http://biorxiv.org/lookup/doi/10.1101/2023.01.21.524489<x> (2023) doi:10.1101/2023.01.21.524489.

98. Makin, J. G., Moses, D. A. & Chang, E. F. Machine translation of cortical activity to text with an encoder–decoder framework. Nature Neuroscience 23, 575–582 (2020).

99. Markowitz, D. A., Wong, Y. T., Gray, C. M. & Pesaran, B. Optimizing the Decoding of Movement Goals from Local Field Potentials in Macaque Cortex. J. Neurosci. 31, 18412–18422 (2011).

100. Tchoe, Y. et al. Human brain mapping with multithousand-channel PtNRGrids resolves spatiotemporal dynamics. Sci. Transl. Med. 14, eabj1441 (2022).

101. Rachinskiy, I. et al. High-Density, Actively Multiplexed µECoG Array on Reinforced Silicone Substrate. Front. Nanotechnol. 4, 837328 (2022).

102. Shi, Z. et al. Silk-Enabled Conformal Multifunctional Bioelectronics for Investigation of Spatiotemporal Epileptiform Activities and Multimodal Neural Encoding/Decoding. Adv. Sci. 6, 1801617 (2019).

103. Zhou, Y. et al. A silk-based self-adaptive flexible opto-electro neural probe. Microsyst Nanoeng 8, 118 (2022).

104. Pesaran, B., Pezaris, J. S., Sahani, M., Mitra, P. P. & Andersen, R. A. Temporal structure in neuronal activity during working memory in macaque parietal cortex. Nat Neurosci 5, 805–811 (2002).

105. Andersen, R. A., Musallam, S. & Pesaran, B. Selecting the signals for a brain–machine interface. Current Opinion in Neurobiology 14, 720–726 (2004).

106. Scherberger, H., Jarvis, M. R. & Andersen, R. A. Cortical Local Field Potential Encodes Movement Intentions in the Posterior Parietal Cortex. Neuron 46, 347–354 (2005).

107. Jun Zhuang, Truccolo, W., Vargas-Irwin, C. & Donoghue, J. P. Decoding 3-D Reach and Grasp Kinematics From High-Frequency Local Field Potentials in Primate Primary Motor Cortex. IEEE Trans. Biomed. Eng. 57, 1774–1784 (2010).

108. Bansal, A. K., Truccolo, W., Vargas-Irwin, C. E. & Donoghue, J. P. Decoding 3D reach and grasp from hybrid signals in motor and premotor cortices: spikes, multiunit activity, and local field potentials. Journal of Neurophysiology 107, 1337–1355 (2012).

109. Bansal, A. K., Vargas-Irwin, C. E., Truccolo, W. & Donoghue, J. P. Relationships among low-frequency local field potentials, spiking activity, and three-dimensional reach and grasp kinematics in primary motor and ventral premotor cortices. Journal of Neurophysiology 105, 1603–1619 (2011).

